# Conformational states of the microtubule nucleator, the γ-tubulin ring complex

**DOI:** 10.1101/2023.12.19.572162

**Authors:** Brianna Romer, Sophie M. Travis, Brian P. Mahon, Collin T. McManus, Philip D. Jeffrey, Nicolas Coudray, Rishwanth Raghu, Michael J. Rale, Ellen D. Zhong, Gira Bhabha, Sabine Petry

## Abstract

Microtubules (MTs) perform essential functions in the cell, and it is critical that they are made at the correct cellular location and cell cycle stage. This nucleation process is catalyzed by the γ-tubulin ring complex (γ-TuRC), a cone-shaped protein complex composed of over 30 subunits. Despite recent insight into the structure of vertebrate γ-TuRC, which shows that its diameter is wider than that of a MT, and that it exhibits little of the symmetry expected for an ideal MT template, the question of how γ-TuRC achieves MT nucleation remains open. Here, we utilized single particle cryo-EM to identify two conformations of γ-TuRC. The helix composed of 14 γ-tubulins at the top of the γ-TuRC cone undergoes substantial deformation, which is predominantly driven by bending of the hinge between the GRIP1 and GRIP2 domains of the γ-tubulin complex proteins. However, surprisingly, this deformation does not remove the inherent asymmetry of γ-TuRC. To further investigate the role of γ-TuRC conformational change, we used cryo electron-tomography (cryo-ET) to obtain a 3D reconstruction of γ-TuRC bound to a nucleated MT, providing insight into the post-nucleation state. Rigid-body fitting of our cryo-EM structures into this reconstruction suggests that the MT lattice is nucleated by spokes 2 through 14 of the γ-tubulin helix, which entails spokes 13 and 14 becoming more structured than what is observed in apo γ-TuRC. Together, our results allow us to propose a model for conformational changes in γ-TuRC and how these may facilitate MT formation in a cell.

## INTRODUCTION

Microtubules (MTs) are dynamic polymers that are formed by head-to-tail protofilaments of α/β-tubulin dimers assembled into a hollow cylinder. MTs are found in all eukaryotes and function in many diverse and essential cellular processes, such as organelle positioning, intracellular transport, and, during cell division, the segregation of chromosomes. For all these critical functions, it is imperative that MTs are generated, or nucleated, at the correct place and time. While spontaneous nucleation of MTs can be driven by high concentrations of tubulin dimers in vitro, nucleation of new MTs in the cell is tightly controlled and requires catalysis^1, 2^. Essential for cellular MT nucleation is the universal MT template, a 2.2 MDa complex known as the γ-tubulin ring complex (γ-TuRC)^3, 4^.

γ-TuRC was first discovered and characterized in *Xenopus laevis* meiotic egg extract^5^, where it was shown to form a cone-shaped structure capable of both nucleating and capping MTs. However, much of the initial work on γ-TuRC focused on the more biochemically and structurally tractable *Saccharomyces cerevisiae* complex. In the cytoplasm, yeast γ-TuRC exists as the so-called γ-tubulin small complex (γ-TuSC), a ‘V’-shaped hetero-tetramer composed of two γ-tubulin molecules bound to the spoke proteins Spc97 and Spc98^6, 7^. When recruited to the MT-nucleating spindle pole body by Spc110, seven γ-TuSCs assemble into a cone topped by a helix of 14 γ-tubulins^8^. This γ-tubulin helix can nucleate MTs in vitro and somewhat mimics the geometry of a 13-protofilament MT, albeit with a wider diameter. Strikingly, artificially “closing” yeast γ-TuRC to match the diameter of a MT—by introducing disulfide-based crosslinks into γ-tubulin—enhances the complex’s nucleation activity three-fold^9^. This result thus prompted the hypothesis that, in the cell, γ-TuRC would change conformation and “close” to a symmetric, MT-like active state during the nucleation reaction.

Recent advances in cryo-electron microscopy (cryo-EM) have led to the determination of the structure of vertebrate γ-TuRC^10, 11, 12^. Like yeast γ-TuRC, vertebrate γ-TuRC is roughly cone-shaped, topped by a 14 γ-tubulin helix. However, vertebrate γ-TuRC contains complexity and asymmetry not present in yeast γ-TuRC. At spoke positions 9-12, GCP2 and GCP3 (the vertebrate orthologs of Spc97 and Spc98, respectively), are replaced by their paralogs GCP4, GCP5, and GCP6. This substitution has been previously reported to cause this region of the γ-tubulin helix to form a less symmetric structure than the γ-TuSC spokes^10^ and thus a poorer match to the highly symmetric MT helix. Furthermore, the base of the cone is stabilized by monomeric actin, the N-termini of GCP3 and GCP6, and small MZT proteins^13, 14^. Likely due to these additional, stabilizing interactions, vertebrate γ-TuRC remains assembled in the cytoplasm^10, 11, 12, 14^, and CDK5RAP2, the vertebrate homolog of Spc110, promotes only γ-TuRC recruitment and activation, not assembly^15, 16^. However, most importantly, vertebrate γ-TuRC also appears to exist predominantly in an inactive, “open” state^10, 11^. How, and even if, vertebrate γ-TuRC changes conformation to an activated, nucleation-competent state remains unclear.

In this work, we have determined structures of two conformations of *X. laevis* γ-TuRC, allowing us to characterize possible conformations of γ-TuRC that may underlie nucleation. We find that the γ-tubulin helix at the top of the γ-TuRC cone is able to undergo substantial deformation due to bending at an internal hinge within individual spokes. However, interestingly, this deformation does not resolve the intrinsic asymmetry observed in vertebrate γ-TuRC. In addition, we have obtained a 3D reconstruction of the γ-TuRC capped MT via cryo-electron tomography, allowing us to visualize a low-resolution envelope of the complex. Our structures combined with the cryo-ET data allow us to propose a model for how conformational changes within γ-TuRC may lead to the generation of MTs.

## RESULTS

### Activity of purified *Xenopus laevis* γ-TuRC

In order to investigate the structure of γ-TuRC, we obtained γ-TuRC from *Xenopus laevis* meiotic egg extract using an endogenous purification strategy^15^, which takes advantage of the high-affinity interaction between γ-TuRC and a fragment of CDK5RAP2 known the γ-TuNA (γ-TuRC nucleation activator). Next, we used a single-molecule MT nucleation assay to assess the activity of our purified γ-TuRC^12, 15^. Coverslips were coated with biotinylated γ-TuRC, then incubated with a MT nucleation mixture containing fluorescently labeled tubulin and either 6 μM γ-TuNA or buffer. MT nucleation from the immobilized γ-TuRC molecules was then monitored via total internal reflection fluorescence (TIRF) microscopy. Consistent with previously published results, we found that our purified γ-TuRC was capable of nucleating MTs (Figure 1a). Additionally, when reactions were supplemented with exogenous γ-TuNA activator, the rate of MT nucleation increased 6.7-fold (Figure 1b)^15^, while other reaction parameters remained unchanged (Supplementary Figure 1a,b). Taken together, these results confirmed the suitability of our purified γ-TuRC for structural studies.

**Figure 1:**
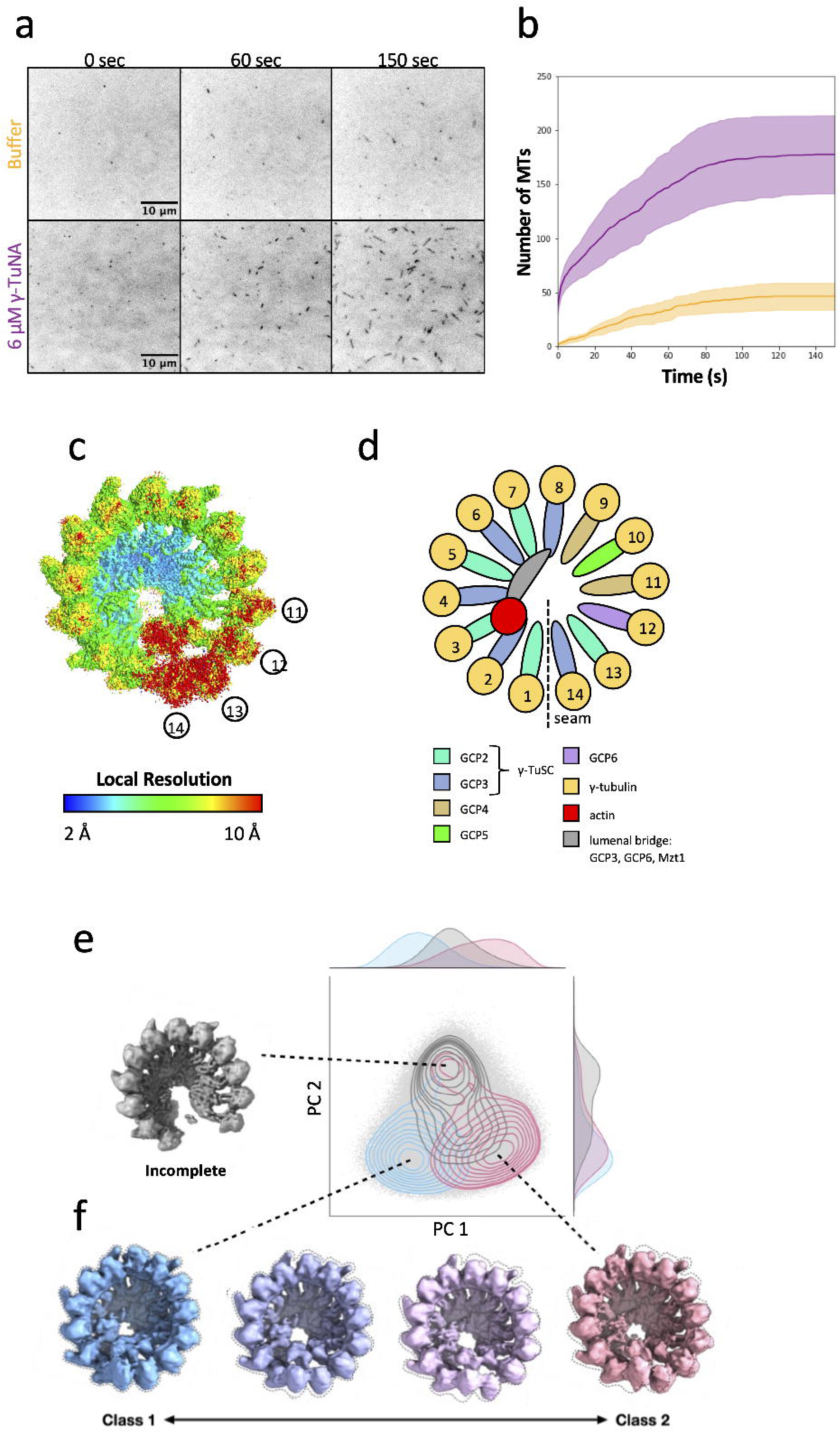
Cryo-EM reconstruction of the conformational landscape of active γ-TuRC. a. Time course of MT nucleation by purified *X. laevis* γ-TuRC. MTs are false-colored dark and appear over time, either in the presence or absence of 6 μM γ-TuNA activating peptide. b. Quantification of MT number over time from images such as the example presented at left. MT nucleation by unactivated γ-TuRC is shown in yellow, and MT nucleation in the presence of activator is shown in purple. Line is shown as the mean of 3 independent reactions, and shaded areas are the standard error of the mean. c. Single particle consensus map of *X. laevis* γ-TuRC, colored by local resolution, where high resolution areas are blue and low resolution areas are colored red. Selected spoke numbers are indicated to orient the reader to areas of differing resolution. d. Cartoon of the structure of γ-TuRC (left), displaying spokes color-coded by the identity of the GCP that forms them. γ-TuRC is displayed here, as always, with the seam or overlap between spoke 1 and spoke 14 facing down. Below, a legend is given identifying key subunits of γ-TuRC according to the color they are displayed as throughout the manuscript. e. The cryoDRGN volume PC space is shown (light gray dots), overlaid with a density plot of particles with associated class labels from the 3DVA analysis. The clustering suggests two main clusters along the vertical axis. The top cluster (gray contour) corresponds to “incomplete” particles with disordered spoke 11–14 density and the bottom to intact particles. The bottom cluster can be broken into two subclusters along the horizontal axis, corresponding to 3DVA classes 1 (blue contour, left) and 2 (pink contour, right). A representative “incomplete” volume map from cryoDRGN is shown in gray at the top left. f. 4 volumes within the 3DVA trajectory are shown, displaying the transition between the Class 1 (blue) and Class 2 (pink) volumes.

### Cryo-EM reconstruction of *Xenopus laevis* γ-TuRC

Next, we proceeded to structural determination of γ-TuRC. Because we wanted γ-TuRC to be in its most active state, we adapted the conditions used for MT nucleation, supplementing the γ-TuRC sample with 6 μM γ-TuNA. We then collected a large single-particle cryo-EM dataset and performed extensive 3D classification and refinement in cryoSPARC^17^ (Supplementary Table 1, Supplementary Figure 2 and 3), resulting in an cryo-EM density map of γ-TuRC at a global resolution of 3.1 Å (Figure 1c, Supplementary Figure 4). In line with previous cryo-EM analysis of γ-TuRC^10, 11, 12^, different regions of the complex displayed substantially different local resolution estimates. The base of the γ-TuRC cone reached the highest resolutions, and the top of the cone, corresponding to the γ-tubulin helix, had a lower average resolution (Figure 1c,d). Spokes 13-14 had the lowest resolution of the entire complex, suggesting substantial conformational and/or compositional heterogeneity.

### **γ**-TuRC undergoes substantial molecular motion

To examine whether and how particle heterogeneity was contributing to regions of low resolution in our map, we first used 3D variability analysis (3DVA)^18^, a method for calculating the principal components (PCs) of variability from a consensus cryo-EM structure. We analyzed the three largest principal components of variability (Supplementary Videos 1-3), which encompassed clamshell-like motions hinging at different locations around the γ-tubulin helix. Across all these principal components of variability, the base of the γ-TuRC cone moves the least, whereas top, corresponding to the γ-tubulin helix, has the most dramatic motions. From this analysis, we found that γ-TuRC appears to undergo a complex series of conformational changes, and, moreover, that these motions entail substantial re-arrangement of the γ-tubulin helix.

To ensure that we were capturing the full range of conformational change, and as a complementary analysis, we explored conformational variability within our data using cryoDRGN^19^, a deep learning algorithm that can extract complex, non-linear molecular motions from single-particle cryo-EM data. CryoDRGN found two major clusters representing compositional and conformational heterogeneity: an outlier cluster (top, gray) that lacked density for spokes 13 and 14, thus representing incomplete complexes, and a second cluster corresponding to intact γ-TuRC particles (Figure 1e, Supplementary Video 4). Reconstructions sampled from the second, intact particle cluster exhibited conformational motions identical to the largest component of variability from 3DVA, and the conformational extremes from 3DVA could be visualized in the space as distinct regions of the latent space (Figure 1f).

### Molecular models of two major **γ**-TuRC conformations

To understand the molecular basis of this conformational change in γ-TuRC, we reconstructed volumes corresponding to the two conformational extremes of the bottom, intact particle cluster, which we will refer to as Class 1 and Class 2 (Figure 1c, Supplementary Video 5), and obtained refined density maps of similar quality for these two classes (Supplementary Figure 5-7). Next, we proceeded to build molecular models for both classes, starting from previously published structures^10, 11, 13, 14^ and performed model refinement (Table 1).

Based on the resolution of our Class 1 and Class 2 maps, we were able to model regions that have thus far remained unresolved, adding to the completeness of prior γ-TuRC models. We observed peptide density on spokes 1 and 2 in the γ-TuRC lumen that was consistent in sequence with the N-terminus of GCP6 (Supplementary Figure 8a). This result suggests an even greater role of GCP6 in γ-TuRC assembly than previously reported^14^, as GCP6 comes into contact with every spoke in γ-TuRC except for spoke 14. We additionally scrutinized the outside surface of γ-TuRC for density corresponding to the γ-TuNA activating peptide. The first potential location for γ-TuNA reported was an extended peptide region termed the “staple” that was found in human γ-TuRC between each GCP2 and GCP3 of a γ-TuSC pair^10^. This density was also visible in our structure; however, in agreement with the more recent structure of recombinant human γ-TuRC ^14^, we interpreted this region as an N-terminal stretch of GCP2 (Supplementary Figure 8b). An updated model of human γ-TuRC bound to γ-TuNA^13^ instead attributed coiled-coil density on the outside of spoke 13 to the γ-TuNA dimer, held in place by the N-terminus of the spoke 13 GCP2 bound to MZT2. Despite extensive refinement of spoke 13, in neither the Class 1 nor the Class 2 map could we observe density consistent with γ-TuNA nor the GCP2/MZT2 module on that spoke (Supplementary Figure 8c). We also did not observe density consistent with γ-TuNA at any previously unattributed position. We suspect the absence of clear γ-TuNA density may be attributed to a difference in binding conditions or else to an intrinsic difference in binding affinity and/or location between human and *Xenopus* γ-TuRC.

### Global rearrangement of **γ**-tubulin subunits during conformational change

The prevailing model of γ-TuRC activation predicts that γ-TuRC will be able to change conformation to better match the geometry of a MT. To date in the field, this global conformational change has been described in terms of changes in the diameter of the γ-tubulin helix^10, 11^. However, our analysis was confounded when we used this metric to compare our two conformations, as we found that the radius to the helical axis varied substantially depending on which spoke of the ring we examined (Figure 2a,b). At certain locations within the ring, centered on spokes 1 and 10, the radius was the narrowest and approached MT radius (Figure 2b, dashed line); however, the ring buckled out at spokes 7-8 for Class 1, 5-6 for Class 2, and 13-14 for both classes. In addition, measuring radius alone did not allow us to capture other ways that the γ-tubulin helix might change conformation, such as alterations in helical pitch or helical rise (Figure 2c). Thus, we developed a new and more complete approach to capture γ-tubulin displacement. First, we extracted the displacement vector required to move each γ-tubulin from its position in Class 1 to its position in Class 2 (Figure 2c,d). Then, we extracted three orthogonal directions of motion—left/right, down/up, and in/out—that describe how each γ-tubulin moves in the frame of reference of the γ-tubulin helix (Figure 2d).

**Figure 2:**
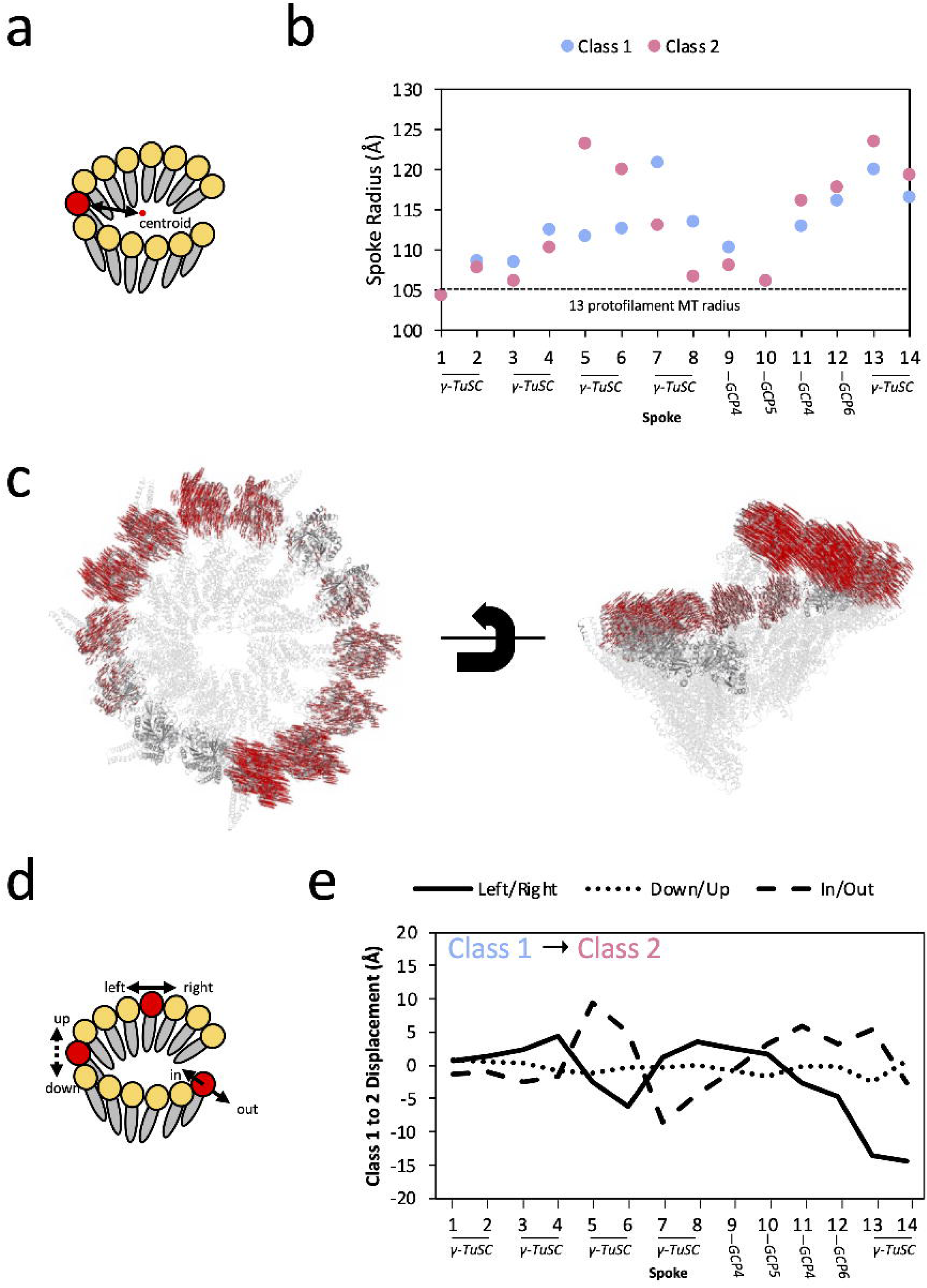
Global Conformational Change within γ-TuRC. a. Cartoon for calculating the radius of γ-TuRC at each spoke from its γ-tubulin (red) to the axis of the γ-tubulin helix (black dashed line). b. Radius of γ-TuRC measured for Class 1 (blue) and Class 2 (pink) at each of the 14 spokes. The radius was measured from the center of each γ-tubulin subunit (approximated by the Cα of Tyr-169) to the central axis of the γ-tubulin helix. As a point of comparison, the radius of tubulin in the 13 protofilament MT, taken as half the average between the outer diameter of 250 Å and inner diameter of 170 Å, is shown as a dashed line at 105 Å. c. All Cα displacement vectors for γ-tubulin residues between Class 1 and Class 2. Class 1 of γ-TuRC is shown as a gray cartoon, where non-γ-tubulin components are partially transparent. Displacement vectors required to move each γ-tubulin Cα to its position in Class 2 are shown as red lines. d. Schematic for measuring orthogonal displacement vectors of each γ-tubulin subunit between Class 1 and Class 2. The overall displacement vector of the Tyr-169 centroid of each γ-tubulin between the two conformations was decomposed into three orthogonal components: left/right, along the vector between γ-tubulin residues Ile-318 to Arg-124, down-up, along the vector between Arg-244 to Asn-102, and in-out, along the vector between Ser-226 to Asp-159. e. Measurement of the displacements of the 14 γ-tubulins of γ-TuRC from Class 1 to Class 2, calculated using the strategy in panel (C). Three components of motion are plotted: left (negative) to right (positive), as a solid line; down (negative) to up (positive), as a dotted line, and in (negative) to out (positive), as a dashed line.

By comparing all three types of motion across all 14 tubulins of the ring, a few major trends emerge (Figure 2e). First, most of the displacement occurs in the left/right and in/out directions, and very little displacement of γ-tubulin up or down in the plane of the helix is observed. Secondly, displacement in these two orthogonal directions is coupled. Displacement into the ring (ring closure, dashed line in Figure 2e) likely results in more crowding in the γ-TuRC lumen, and thus is generally coupled with a clockwise rotation towards spoke 14 (unspooling of the γ-tubulin helix, solid line in Figure 2e). To counteract this global deformation, displacement out of the ring (ring opening, dashed line) is, conversely, coupled with a counterclockwise rotation towards spoke 1 (respooling of the γ-tubulin helix, solid line). Many parts of the ring, most notably spokes 9-12, appear to be relatively static between the two conformations (Supplementary Figure 9), and the largest conformational difference between Class 1 and Class 2 appeared to be an exchange in two γ-TuSCs—in Class 1, the spoke 5-6 γ-TuSC leans into the ring and the spoke 7-8 γ-TuSC leans out, whereas in Class 2, the spoke 5-6 γ-TuSC now leans out of the ring and the spoke 7-8 γ-TuSC leans in. In conclusion, any “closure” of the γ-tubulin helix in a subset of spokes appears to be coupled with an equivalent “opening” elsewhere in the ring.

### Local asymmetry of the **γ**-tubulin helix is preserved during conformational change

For activated γ-TuRC to become a more perfect MT template, the γ-tubulin helix will need to match the geometry of the MT not only in terms of global parameters—for example, diameter— but also locally. In particular, the spacing between adjacent γ-tubulins will need to match the spacing between MT protofilaments. Therefore, we measured the distance between γ-tubulin subunits in adjacent spokes and compared with the distance between protofilaments in the MT helix (Figure 3a). In contrast to the global displacement of γ-tubulins of up to 10-15 Å between Class 1 and Class 2 described above (Figure 2e), the two classes had similar separation between adjacent γ-tubulins (Figure 3b, Supplementary Figure 10a). The largest distance change, 3 Å, was located between spokes 12 and 13; however, due to the low local resolution in this region, we cannot conclude that this change reflects a true difference between the two classes.

**Figure 3:**
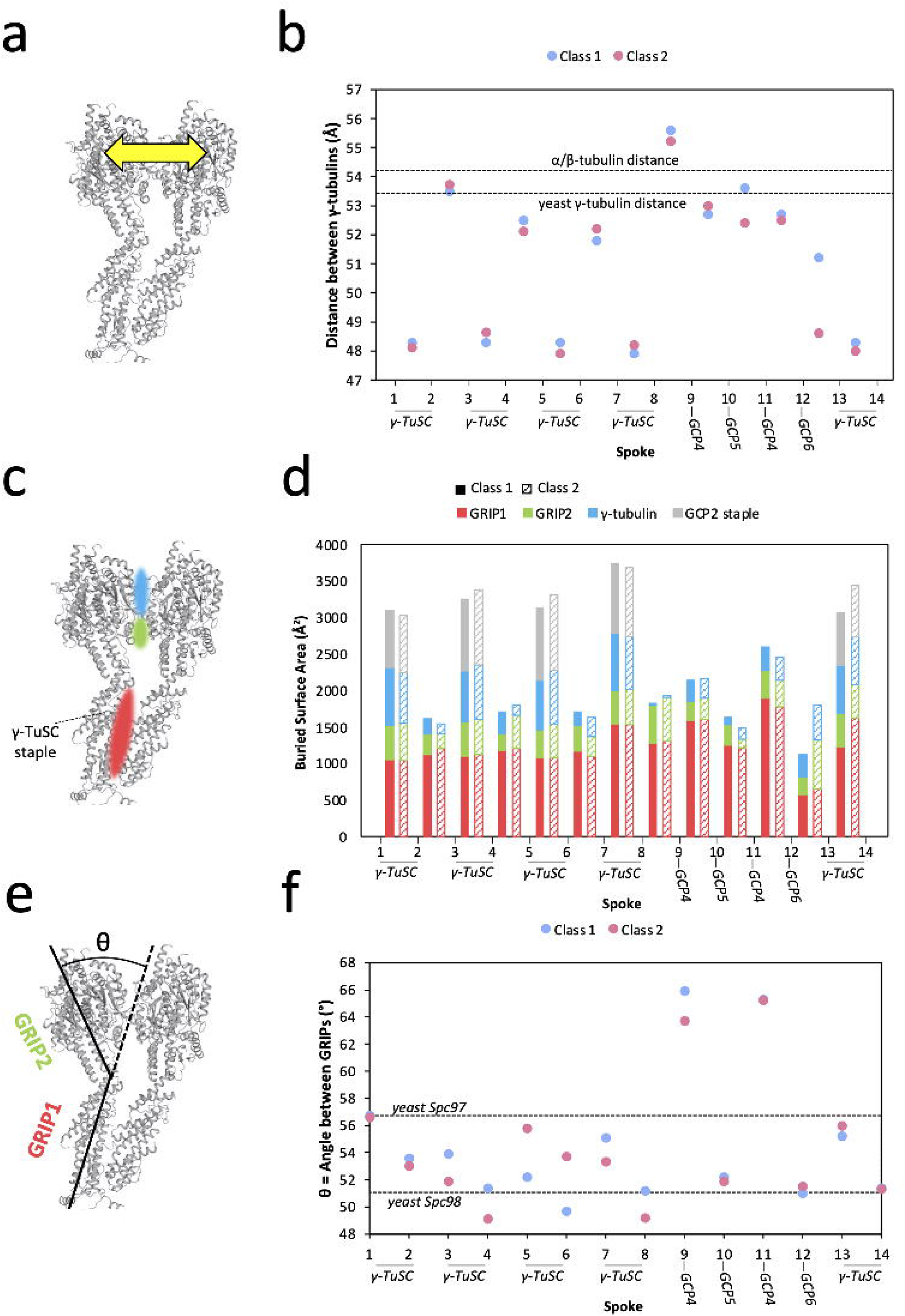
Local asymmetry of γ-TuRC leads to template imperfection. a. Schematic for measuring distances between adjacent γ-tubulin subunits. Distances were measured from the centroid residue Tyr-169. b. Measurement of distances between adjacent γ-tubulin subunits. Distances for Class 1 are shown in blue and distance for Class 2 in pink. Measurements were calculated as schematized in (A). Dashed lines indicate reference distances between adjacent γ-tubulin subunits in the closed yeast γ-TuRC (PDB accession 7M2W), measured between the γ-tubulin centroid residue Tyr-170 as 53.6 Å around the symmetric helix, and between adjacent β-tubulin subunits in the 13 protofilament MT (PDB accession 7SJ7), measured between the β-tubulin centroid residue Phe-167 as 54.2 Å around the symmetric helix. c. Schematic for measuring buried surface areas between adjacent γ-TuRC spokes. Colored patches represent contact areas where buried surface area is measured. Red represents GRIP1, the N-terminal half of the GCP, green represents GRIP2, the C-terminal half of the GCP, and blue represents γ-tubulin. d. Measurement of buried surface area between adjacent γ-TuRC spokes. Buried surface area is tabulated for GRIP1 (red), GRIP2 (green), and γ-tubulin (blue). When the contact is between GCP2 and GCP3 within a γ-TuSC, the surface area buried by the staple is also tabulated. Surface areas for Class 1 are shown as solid bars and, for Class 2, as striped bars. The precise residues corresponding to GRIP domain assignments can be found in the Methods section. e. Schematic for measuring the angle between the GRIP1 and GRIP2 domains of GCP subunits. f. Measurements of the angle between GRIP1 and GRIP2 domains for each spoke of Class 1 (blue) and Class 2 (pink). Dashed lines show the angle displayed for yeast Spc97 (analogous to GCP2, 56.4°, and yeast Spc98 (analogous to GCP3, 51° in the closed yeast γ-TuRC (PDB accession 7M2W). Residues used to calculate domain angles can be found in the Methods section.

While any difference between Class 1 and Class 2 inter-γ-tubulin distances was generally less than 1 Å, the variation in distance between γ-tubulins of adjacent spokes around the ring was much more pronounced, up to 6 Å, independent of the class (Figure 3b). In the region of γ-TuRC composed of γ-TuSCs, γ-tubulin distances alternate: within a γ-TuSC pair (e.g. spoke 1 to 2) γ-tubulins are closely spaced, whereas between γ-TuSCs (e.g. spoke 2 to 3), γ-tubulins are spaced further apart (as also noted in ^11^). However, where GCP4/5/4/6 substitutes for γ-TuSCs, this pattern is broken, and all γ-tubulins are separated by the same, larger spacing (Figure 3b).

We next wanted to understand how γ-tubulin spacing could relate to ring closure; therefore, we compared these distances to those found between the protofilaments of a MT (dashed line in Figure 3b). In the MT, tubulin dimers are separated by 54.2 Å (PDB accession 7SJ7^20^), and, surprisingly, this distance is most consistent with the further apart, or more open distances in γ-TuRC (Figure 3b). This is also the distance found between γ-tubulins in the “closed” disulfide-linked active yeast γ-TuRC (PDB accession 7M2W^21^). Therefore, the most “closed,” or closest, γ-tubulin distances within vertebrate γ-TuRC deviate the most from MT geometry and appear to be too closely spaced to properly nucleate a MT.

Is γ-TuRC likely to be able to change conformations to “open” its γ-TuSC pairs to a MT-like geometry? To answer this question, we investigated the intermolecular contacts that drive association of adjacent γ-TuRC spokes. By comparing buried surface area (BSA) (Figure 3c), we found that BSA changed little between Class 1 and Class 2, but that spokes that comprised a γ-TuSC pair had greater BSA than other spoke pairs (Figure 3d, Supplementary Figure 10b-d). This increase in surface area results both from increased contact between γ-tubulins, as well as from the presence of the staple peptide sandwiched between GCP2 and GCP3, and it suggests that each γ-TuSC is a relatively rigid, autonomous unit. This result is in line with prior work demonstrating that pre-assembled γ-TuSC units load onto the assembling γ-TuRC^11, 14^; however, it also suggests that separating the γ-tubulins of a vertebrate γ-TuSC to resemble the MT would be energetically unfavorable.

### Role of GCPs in positioning the **γ**-tubulin helix

If neither the distances between γ-tubulins nor the areas buried between adjacent spokes change substantially between Class 1 and Class 2, what then leads to the rearrangement we observe within the γ-tubulin helix? The most substantial inter-spoke contacts in γ-TuRC are found at the base of the cone, within the GRIP1 domain of the GCPs, and, in contrast, GRIP2 and γ-tubulin generate relatively minor and variable contact surfaces. Therefore, we wondered whether a hinge movement between the two GRIP domains of each spoke could lead to γ-tubulin repositioning (Figure 3e). When we measured the angle between GRIPs, we found that it varied up to 7° depending on the spoke measured and up to 4° between Class 1 and Class 2 (Figure 3f). As previously described, the largest conformational changes between Class 1 and Class 2 represent a “flip-flop” exchange of spokes 5-6 and spokes 7-8 (Figure 2b,e), and these spokes also represent the largest changes in GRIP angle between Class 1 and Class 2 (Figure 3f). More specifically, increase in the angle between GRIP1 and GRIP2 correlates with displacement of a spoke out of the ring. Consistent with this general pattern, the GRIP1/2 angle of spokes 5-6 is greater in Class 2 than Class 1 (Figure 3f), and in Class 2 these spokes lean out of the ring in comparison with Class 1. The opposite pattern is observed for spoke 7-8, which lean into the ring in Class 2 when compared with Class 1. Thus, bending of the GRIP1/2 hinge appears to be linked to the conformational change observed between the two γ-TuRC classes. In contrast to γ-tubulin distances, we find that the range of angles adopted by vertebrate GRIPs is similar to that found in yeast γ-TuRC, with the exception of GCP4^11^. However, GRIP angles are far more asymmetric around the vertebrate ring than are their yeast counterparts.

### Cryo-ET of a **γ**-TuRC nucleated and capped MT

Up to this point, we have examined spontaneous conformational change within γ-TuRC, which may not correlate directly with the changes that occur during MT nucleation. To gain a better understanding of MT nucleation, we turned to cryo-electron tomography (cryo-ET), with the goal of studying γ-TuRC in complex with MTs. Because MT nucleation and MT polymerization compete for the same starting material (namely, soluble tubulin dimers), in order to maximize MT nucleation complexes, we needed to also suppress MT polymerization. To accomplish this task, an engineered MT capping protein, known as DARPin TM-3^22, 23^, was titrated into the reaction at a concentration that restricts MT length to about 1 μm (Figure 4a, Supplementary Figure 11). After MTs capped with γ-TuRC were nucleated in the presence of DARPin TM-3 and the MT stabilizer paclitaxel, 105 tilt series were collected and reconstructed into tomograms (Supplementary Table 1). In the resulting tomograms, we were able to easily differentiate γ-TuRC capped MTs from open MT ends (Figure 4b).

**Figure 4:**
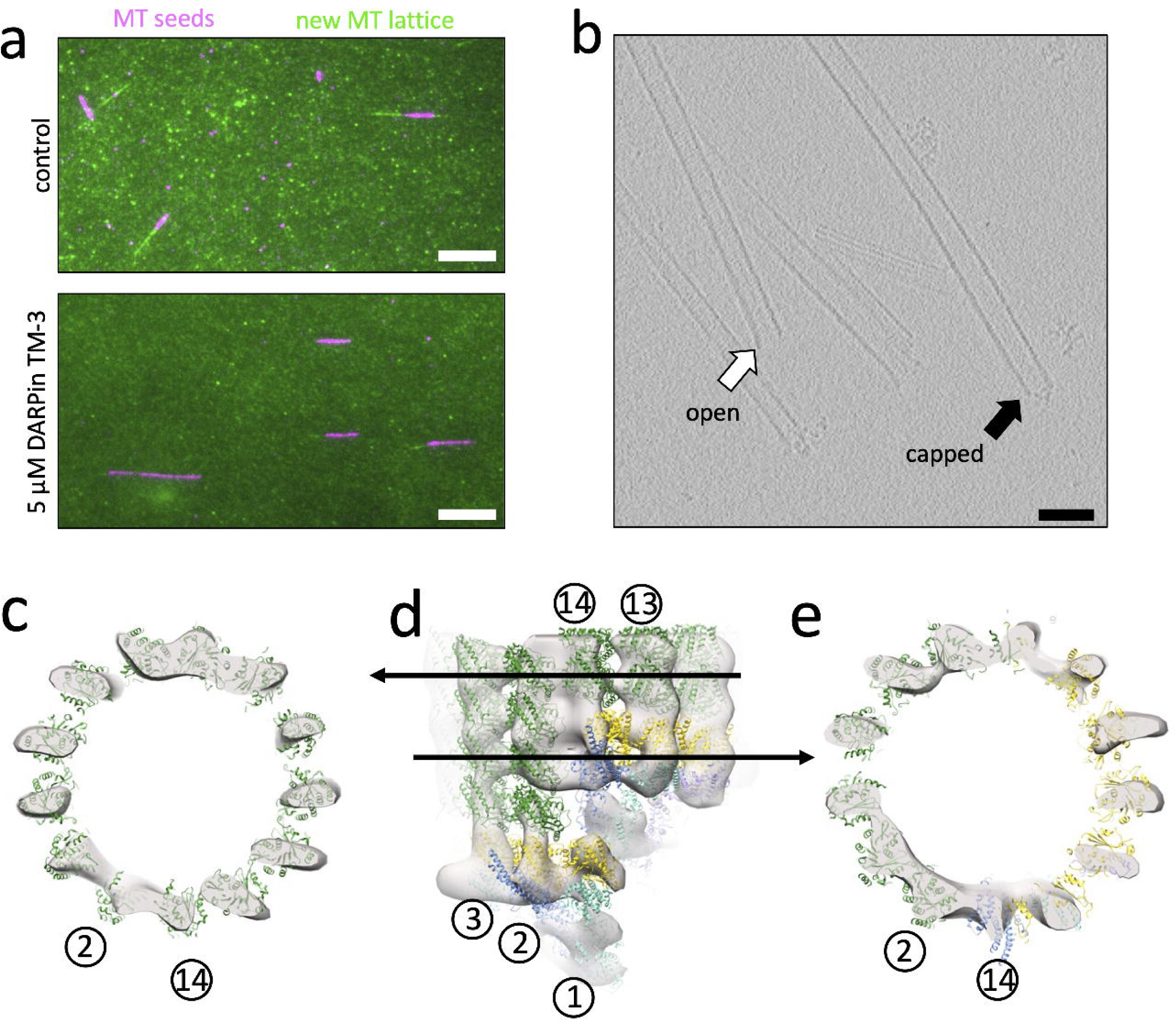
Cryo-ET reconstruction of the γ-TuRC capped MT minus end. a. Representative still images from MT growth assays, where pre-formed, GMPCPP-stabilized and Atto-647 labeled MT seeds (purple) were affixed to coverslips, and the polymerization speed of new MT lattice was measured by monitoring addition of Alexa-488 labeled soluble tubulin (green). Polymerization reactions occurred in the absence (top) or presence (bottom) of the plus-end capping protein DARPin TM-3 at 5 μM and were measured at a time point of 5 minutes. Scale bar is 10 μm. b. Representative slice from a tomogram showing both open (spontaneously nucleated, white arrow), and capped (γ-TuRC bound and nucleated, black arrow) MTs. Scale bar is 10 nm. c. Top view of a slice through the MT-only region of the cryo-ET reconstruction of the capped MT (transparent gray surface; here and in the remainder of the figure, low-pass filtered to 34 Å). A model of a 13-protofilament MT (PDB accession 7SJ7), where α-tubulin is shown in dark green and β-tubulin in light green is docked into the cryo-ET density, showing the position of the 13 protofilaments. For reference as to the orientation of γ-TuRC within the cryo-ET reconstruction, the position of spokes 2 and 14 is given as reference (circled numbers). Note that this orientation is slightly offset from those shown elsewhere in the manuscript, as the axis of γ-TuRC and the axis of the MT do not fully coincide. d. Side view of the cryo-ET reconstruction of the γ-TuRC-capped MT, with Class 1 γ-TuRC, colored as elsewhere in the figure, docked into the cone-shaped density and the docked MT above it. Arrows indicate the location within the cryo-ET reconstruction of the slices shown in panels A and C. e. Top view of a slice through the mixed α-tubulin/γ-tubulin helix of the MT template. The 13-protofilament pattern extends around the ring; however, protofilaments at spokes 9-13 are composed of γ-tubulin template (yellow), whereas protofilaments at spokes 2-8 are composed of α-tubulin MT (dark green).

After subtomogram averaging of 94 γ-TuRC-capped MT volumes, we were able to generate a reconstruction of the post-nucleation state of γ-TuRC at a low resolution of approximately 35 Å (Figure 4c-e). Whereas in vitro, untemplated MTs have protofilament numbers varying from 11-16, the reconstruction of the γ-TuRC nucleated MT shows an exclusively 13-protofilament MT capped by a cone-shaped structure (Figure 4c). Asymmetric features of γ-TuRC were visible in the cone-shaped density, such as the groove separating spoke 1 and spoke 14 of γ-TuRC at the seam (Supplementary Figure 13a), as well as connecting density between spokes 2 and 8 corresponding to the lumenal bridge structure (Supplementary Figure 13b).

What is the conformation that γ-TuRC adopts after nucleating a MT? Although the resolution of our reconstruction does not allow us to differentiate between the conformations of Class 1 and Class 2, we can gain preliminary insight into the conformational changes resulting in the post-nucleation state of γ-TuRC. Spokes 13 and 14 are highly disordered and apparently prone to dissociation in the apo γ-TuRC structure (Figure 1c, Supplementary Figure 14a). Our initial hypothesis for this disorder was that, during nucleation, spoke 14 needs to move out of the way to uncover the better-positioned spoke 1 to allow spoke 1 to form a portion of the MT template. In this case, disorder of spokes 13-14 would in fact promote nucleation, and the nucleated MT would bind to spokes 1-12. However, our cryo-ET reconstruction was inconsistent with this hypothesis. Instead, the nucleated MT binds to the γ-tubulins of spokes 2-14, while spoke 1 is occluded by spoke 14 (Figure 4d,e). Moreover, in the cryo-ET reconstruction, the GCPs of spokes 13 and 14 appear to have the same amount of density in the reconstruction as the well-ordered spokes 1 and 2 (Figure 4d, Supplementary Figure 14b). This suggests that, post-nucleation, spokes 13 and 14 become more ordered than in the Class 1 or Class 2 apo structure.

Thus, after nucleating a MT, γ-TuRC retains many of its asymmetric features. In addition, despite the mismatches between γ-TuRC and the MT geometry noted above, the MTs nucleated by γ-TuRC are in fact 13 protofilament MTs, the biologically relevant MT geometry. Moreover, MT nucleation and/or MT binding in fact leads to visible differences between post-nucleation γ-TuRC and the apo structures, where spokes 13 and 14 participate in the MT template and, instead of dissociating fully, appear to become more ordered.

## DISCUSSION

How γ-TuRC nucleates MTs has been a major question since the discovery of the complex in 1995^5^. Determination of the structure of the vertebrate complex was a major landmark. An open question is how γ-TuRC changes conformation, particularly as part of its catalytic cycle. Although numerous structures of γ-TuRC have been solved since^10, 11, 12, 13, 14^, the different strategies used for cryo-EM data refinement has made it challenging to compare conformational states across different structures. In this work, we have filled this gap by determining the structure of two conformational states within the same active γ-TuRC sample via single particle cryo-EM. The characterization of these structures allow us to begin to understand some of the patterns underlying motion within γ-TuRC. We have also determined the structure of γ-TuRC in its post-nucleation state, bound to the MT. Taken together, this collection of γ-TuRC structures allows us to address a hypothesis of γ-TuRC nucleation: that, in its inactive, cytoplasmic state, the diameter of γ-TuRC is too large to match the one of a MT and must “close” during activation.

The best experimental evidence for the “ring closure” hypothesis^8^ was the discovery that yeast γ-TuSC adopts two conformational states, one of which represents an active ‘closed’ conformation that was engineered via disulfide links^9^. To date, it still remains to be formally determined whether yeast or other γ-TuRCs adopt a ‘closed’ state during nucleation. However, we can now ask whether vertebrate γ-TuRC is, in fact, even able to undergo ring closure. One of the primary differences between the yeast and vertebrate complexes is that the vertebrate complex has three additional spoke proteins, GCP4/5/6. Thus, the vertebrate complex is inherently more asymmetric than the yeast γ-TuRC. Surprisingly, though, the vertebrate-only subunits GCP4/5/6 form the region of vertebrate γ-TuRC that is the most static and congruent with MT geometry. The yeast-like subunits GCP2 and GCP3, in contrast, represent the region of asymmetry within vertebrate γ-TuRC, and the distance between γ-tubulins within the vertebrate γ-TuSC is substantially shorter than expected. Thus, any activating conformational change is likely to take a much different form between yeast and vertebrate γ-TuRC.

Does vertebrate γ-TuRC need to undergo conformational change in order to nucleate a MT? Previous structures have noted that γ-TuRC lacks the symmetry present in a MT helix^10, 11^. Our work now shows that, despite conformational change, vertebrate γ-TuRC remains an asymmetric, imperfect template within the conformations that we have found. This then raises the question of how this template can nucleate a symmetric end product. Yet, biochemical results with γ-TuRC subcomplexes have demonstrated that asymmetric templates can nucleate MTs. In the most extreme case, linear γ-tubulin oligomers alone are able to nucleate cylindrical MTs^24, 25^. This evidence suggests that a ring, let alone a symmetric one, is not required for MT nucleation. In addition, a four-spoke subcomplex composed of GCP4/5/4/6 can support nucleation at 80% the rate of a 14-spoke intact complex^26^. We find this result significant, because GCP4/5/4/6 is also the most stable and symmetric region within γ-TuRC. Moreover, kinetic results and biophysical modeling from our group have suggested that binding of 4 tubulin dimers to g-TuRC forms a sufficient nucleus to generate a MT^24^. Thus, we propose that the first four tubulin dimers required for the rate-limiting step within nucleation are assembled onto these spokes, thereby facilitating their lateral interactions that are challenging to form in solution.

How might MT nucleation proceed from this 4-spoke nucleus? Our cryo-ET reconstruction further explains the somewhat puzzling observation that tubulin itself can activate nucleation from γ-TuRC^24^. In fact, one of the most glaring imperfections of the apo γ-TuRC as a MT template is the fact that two of its spokes, 13 and 14, are highly disordered. Yet, we find that these spokes do in fact form part of the MT template, and this disorder seems to be reduced upon MT binding. Thus, binding of tubulin to spokes 13 and 14 will in fact likely improve this critical region of the template, allowing extension of the nascent MT from spoke 12. We also hypothesize that γ-TuNA, if it in fact binds to the spoke 12-13 interface as seen in human γ-TuRC^13^, may facilitate nucleation by aiding the positioning of spoke 13 and aiding in lattice extension onto this spoke.

The example of yeast γ-TuRC demonstrates that it may be possible for the cell to form a symmetric 13-spoke template. So, why would vertebrate γ-TuRC form such an asymmetric, imperfect template? We speculate that this intrinsic imperfection of γ-TuRC may in fact be a feature, not a bug, requiring the presence of additional co-nucleating factors and other regulators. This would allow the cell to further control the nucleation activity of γ-TuRC spatiotemporally. The complexity of γ-TuRC’s asymmetry would allow for multiple, conformationally-distinct binding sites for activators and regulators, providing multiple inputs of regulation along the 14 spokes. Future work will be needed to determine how additional γ-TuRC binding partners may alter its conformational dynamics and the overarching structure of the nucleation site to fine-tune nucleation. The activity and structure of this multimodal regulation will be a subject of future studies.

## METHODS

### Protein cloning, expression, and purification

#### DARPin TM-3

The published DARPin TM-3 sequence^22^ was synthesized by GenScript and cloned into a pST50 expression vector^27^ with N-terminal 6x His tag using Gibson assembly. The plasmid was transformed into Rosetta2 (DE3) pLysS cells (Millipore). Rosetta2 cells were then grown in 4 L of Luria Broth Miller (LB) at 37°C to an optical density of 0.6, then expression was induced with 0.5 mM isopropyl-β-D-thiogalactopyranoside (IPTG) at 16°C for 18 hours. The cells were pelleted, snap frozen in liquid nitrogen, and stored at −80°C.

To purify the TM-3 construct, 1 L of cell pellet was thawed on ice and resuspended into Lysis Buffer (50 mM Tris•HCl, pH 8, 300 mM NaCl, 15 mM imidazole, 6 mM β-mercaptoethanol, 1 μM phenylmethylsulfonyl fluoride (PMSF), 10 μg DNase I from bovine pancreas) and a single dissolved cOmplete EDTA-free Protease Inhibitor Cocktail tablet (Roche). The pellet was resuspended using a Biospec Tissue Tearor (Dremel) and lysed using an Emulsiflex C3 (Avestin) at 10,000-15,000 psi. The cell lysate was spun at 33,000 x *g* for 40 minutes at 4°C.

Affinity purification was performed via a HisTrap HP column (Cytiva), which was previously equilibrated with His Binding Buffer (50 mM Tris•HCl, pH 8, 300 mM NaCl, 15 mM imidazole, 6 mM β-mercaptoethanol, 1 μM PMSF). The column was then washed with high-salt binding buffer (binding buffer with 1 M total NaCl) followed by low-salt buffer (binding buffer with 10 mM total NaCl). Finally, protein was eluted using binding buffer with 150 mM total NaCl and 250 mM total imidazole. The peak fraction was collected for further purification via SEC.

A HiLoad 16/600 Superdex 75 SEC column (Cytiva) was equilibrated with PKM buffer (Ahmed et al 2016) (20 mM PIPES, pH 6.8, 150 mM KCl, 1 mM MgCl_2_, 0.5 mM ethylene glycol-bis-(β-aminoethyl ether)-N,N,N’,N’-tetraacetic acid (EGTA)), with 6 mM β-mercaptoethanol and 1 μM PMSF added. Sample was loaded onto the column, and the peak fractions were pooled and concentrated using a 3 kDa molecular weight cut off (MWCO) spin concentrator (Merck). The concentration of the protein was measured via Bradford assay against a bovine serum albumin (BSA) protein standard, and the purity was assessed using SDS-PAGE and Coomassie stain. Samples were aliquoted and flash frozen in liquid nitrogen before storage at −80°C.

### Human and *Xenopus* γ-TuNA constructs

Purification of human Strep-His-(TEV)-HaloTag-(3C)-γ-TuNA and *Xenopus* Strep-His-(3C)-γ-TuNA has been previously described in^15^ and was performed similarly to the DARPin TM-3 purification protocol above. In summary, the previously-cloned constructs were expressed in Rosetta 2 (DE3) pLysS *E. coli* cells. The cells were grown in 2 L Terrific Broth cultures to an optical density of 0.7 and induced with 0.5 mM IPTG at 16°C for 18 hours. The cultures were pelleted, snap-frozen, and stored at −80°C, and thawed on ice prior to lysis.

After lysis via Emulsiflex C3 and clarification, the Strep-His-(TEV)-HaloTag-(3C)-human γ-TuNA protein was purified using StrepTactin SuperFlow resin (IBA). After elution using 2.5 mM desthiobiotin, protein purity was assessed by SDS-PAGE, and concentration estimates were performed by Bradford assay using a BSA standard curve. For γ-TuRC purification, Halo-tagged human γ-TuNA was flash-frozen and stored at −80°C. *Xenopus* Strep-His-(3C)-γ-TuNA was purified similarly. Yield and purity were assessed via SDS-PAGE gel and Coomassie stain.

### Endogenous *Xenopus* γ-TuRC Purification via Halo-human γ-TuNA pulldown

*Xenopus* γ-TuRC purification was performed as described in^15^. In brief, 15 mg of His-Tev-HaloTag-(3C)-human γ-TuNA was coupled to Halo Magne beads (Promega). *Xenopus laevis* egg extract was prepared by standard methods^28, 29^ and in accordance with the guidance of the Princeton Institutional Animal Care and Use Committee. 4 mL *Xenopus laevis* egg extract was added to the coupled beads and incubated for 2 hours. After 3 washes with CSF-xB 2% sucrose (10 mM HEPES•KOH, pH 7.7, 100 mM KCl, 1 mM MgCl_2_, 0.1 mM CaCl_2_, 5 mM EGTA, 2% (w/w) sucrose), γ-TuNA bound to γ-TuRC was eluted overnight via cleavage with PreScission 3C protease. In the case of γ-TuRC prepped for nucleation reactions under TIRF microscopy, γ-TuRC was biotinylated prior to elution by incubation for 1 hour on ice in CSF-xB 2% (w/w) sucrose supplemented with 40 μM NHS-PEG_4_-biotin (ThermoFisher). The elution was concentrated using a 100 kDa MWCO spin concentrator (Amicon). The concentrated eluate was further purified and buffer exchanged using a 10-50% (w/w) sucrose gradient made with CSF-xB or BRB80 buffer (80 mM PIPES•KOH, pH 6.8, 1 mM MgCl_2_, 1 mM EGTA) supplemented with 20 μM guanosine-5’-triphosphate (GTP) and 10 μg/mL each leupeptin, pepstatin A, and chymostatin at 200,000 x *g* for 3 hours. The gradient was fractionated and the peak γ-TuRC fraction was determined by Western blot using mouse monoclonal anti-γ-tubulin (T6557, Sigma Aldrich) and negative stain EM. The peak fraction of biotinylated γ-TuRC was aliquoted and snap frozen in liquid nitrogen.

### Solution Reactions and Microscopy

#### Single molecule TIRF for γ-TuRC activation by γ-TuNA

For these experiments, we followed previously published protocols which we will describe here in brief^12, 15, 24^.

High Precision 22×22 mm No. 1.5H (Deckglaser) were cleaned by sonication with 3 M NaOH then Piranha solution (2:3 ratio of 30% (w/w) H_2_O_2_ to sulfuric acid) and treated with 3-glycidyloxypropyl trimethoxysilane (GOPTS) (Sigma). GOPTS was then reacted with a mixture of 9 parts α-hydroxy-ω-amino PEG 3000 to 1 part α-biotinyl-ω-amino PEG 3000 by weight (Rapp Polymere). Flow cells for imaging were made using double-sided tape applied to a glass slide and passivated with 2 mg/mL PLL_20_-g[3,5]-PEG_2_ (SuSOS AG). Functionalized coverslips were placed face-down bridging two pieces of double-sided tape.

To perform nucleation reactions, flow cells were first blocked with 5% (w/v) Pluronic F-127 and assay buffer (BRB80 with 30 mM KCl, 1 mM GTP, 0.075% (w/v) methylcellulose (4000 cP), 1% (w/v) D-glucose, 0.02% (w/v) Brij-35, 5 mM β-mercaptoethanol) supplemented with 0.05 mg/ml κ-casein. 0.05 mg/mL NeutrAvidin (ThermoFisher) in κ-casein buffer was then bound to biotin-PEG before flowing in biotinylated γ-TuRC diluted 1:100 in BRB80. Reaction mixes were made with 10 μM bovine tubulin (PurSolutions) containing 5% Alexa 568-labeled bovine tubulin and 1 mg/mL BSA (Sigma) in assay buffer. We then added either Tris control buffer or Strep-His-3C-*Xenopus* γ-TuNA to 6 μM. The reaction mix was precleared by ultracentrifugation at 280,000 x *g* for 10 minutes, and 0.68 mg/mL glucose oxidase (SERVA GmbH) and 0.16 mg/mL catalase (Sigma) were added. Imaging was performed as in our previous work^15, 24^ on a Nikon Ti-E inverted stand with an Apo TIRF 100x oil objective (NA = 1.49) and an Andor iXon DU-897 EM-CCD camera with EM gain set to 300. The objective was heated to 33.5°C using a heated collar (Bioptechs). Imaging began after flow cells incubated on the warm objective for 1 minute.

To measure MT mass over time, we used the ‘Subtract Background’ function in ImageJ with FIJI plugins^30^ with a rolling ball radius of 50 pixels on all frames, then measured mean MT intensity. We then subtracted the mean intensity from the first frame of each timelapse from all subsequent frames and plotted the average mean MT intensity over time for each condition. To quantitate MT dynamics, we cropped each time series to 256×256 pixels for the first 150 sec of imaging and used the StackReg plugin^31^ to correct for translational drift. We next generated kymographs of MTs and processed them to derive whether they were γ-TuRC nucleated, and, if they were, the frame at which nucleation occurred, the MT growth rate, and maximum MT length. Spontaneously nucleated MTs exhibited bidirectional growth, while γ-TuRC nucleated MTs grew in only one direction from a fixed point on the coverslip. To analyze MT nucleation rates, we plotted the mean cumulative curve of MTs nucleated over time for each condition and displayed ±SEM. We fit these curves the following equation:

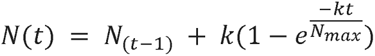

where *N(t*) = the number of MTs nucleated at that time point, *N_max_ =* the maximum number of MTs nucleated after 150 s, and *k* = the γ-TuRC nucleation rate. The average MT growth rate and maximum length for each timelapse was calculated and plotted according to condition. A nonparametric t-test was performed between the two conditions, where p-values less than 0.05 were considered significant.

#### DARPin TM-3 MT polymerization assay and processing

Glass coverslips were first passivated with dichlorodimethylsilane (Sigma), as previously published^32, 33^. These coverslips were then attached with double-sided tape to glass slides to create 2-channel imaging chambers. To each chamber were added 0.2 mg/mL anti-biotin (ab1227, Abcam) in BRB80, followed by 5 mg/mL κ-casein in BRB80 (Sigma Aldrich), and finally washed with BRB80.

Prior to imaging, single-cycled GMPCPP-stabilized, Atto-647 labeled, biotinylated MT seeds were made as previously described^32^ and were first attached to the coverslip. Next, a reaction mixture of 20 mg/mL bovine tubulin, 2 mg/mL Alexa-488 labeled bovine tubulin, 50 mM GTP, and an oxygen scavenging system composed of 2.5 mM protocatechuic acid (PCA), 25 nM protocatechuate-3,4-dioxygenase (PCD), and 2 mM 6-hydroxy-2,5,7,8-tetramethylchroman-2-carboxylic acid (Trolox) in BRB80 with either 100 nM DARPin TM-3 or additional BRB80 was flowed through a chamber. Slides were immediately transferred to the microscope stage.

Slides were imaged on a Nikon Ti-E inverted system, with an Apo TIRF 100x oil objective (NA = 1.49) fitted with an objective heater set to 33.5°C. Images for both conditions (100 μM DARPin and control) were taken in parallel at 10 second intervals for 10 minutes using NIS-Elements AR program (NIKON, ver.5.02.01-Build 1270). with an Andor Neo Zyla (VSC-04209) camera. The MT polymerization movies were processed using ImageJ with FIJI plug-ins^30^. For each condition, both channels were merged into a single movie. Lines were drawn along each MT and the *kymograph* function was used to create a kymograph of each MT.

#### Solution pelleting nucleation reaction with DARPin

A MT polymerization reaction containing ∼100 nM sucrose-gradient purified γ-TuRC, 15 μM pre-cleared bovine brain tubulin, 1 mM GTP, 1.5 μM paclitaxel, 1 mM DTT, 5.25 μM DARPin TM3, and BRB80 buffer was set up on ice. The reaction was then moved to a 37°C water bath for 5 minutes. Upon removal, additional paclitaxel was added to the reaction for a final concentration of 15 μM. The reaction was layered onto a cushion of 40% (w/w) sucrose in BRB80 with 20 μM paclitaxel using a wide-bore pipette tip. MTs were pelleted at 150,000 x *g* for 20 minutes in a Beckman tabletop ultracentrifuge set at 26°C. After the spin, the supernatant was removed, and the pellet resuspended in BRB80 + 20 μM paclitaxel. The sample was left at room temperature and immediately used for grid preparation.

### Grid preparation and EM data collection

#### Preparation of cryo-EM grids of γ-TuRC with γ-TuNA

Prior to grid preparation, purified, cleaved *Xenopus* γ-TuNA was added to BRB80 sucrose-gradient purified γ-TuRC at a final concentration of 6 µM γ-TuNA. The two components were allowed to bind on ice for 20 minutes. A Leica EM GP2 plunge freezer (Leica Microsystems) was set up with the chamber set to 95% humidity and 4°C. Quantifoil R1.2/1.3 400-square mesh Cu grids with 2 nm continuous carbon (Quantifoil) were glow-discharged for 8 s at 10 mA using a PELCO easiGlow glow discharging system (Ted Pella) and loaded into the EM GP2 chamber. 5 μL of sample was added onto the grid, allowed to incubate for 45 seconds, and blotted by hand with Whatman No. 1 blot paper by touching the side of the grid. Then, 3 μL of BRB80 + 0.02% (v/v) Tween 20 was applied to the grid as a wash and front-blotted for 10 seconds using the Leica EM GP2 before plunge-freezing in liquid ethane. The grid was then transferred to a grid box in liquid nitrogen and stored in liquid nitrogen until data collection.

#### Cryo-EM data collection of γ-TuRC with γ-TuNA

Data was collected at NYU Langone Health Cryo-Electron Microscopy Laboratory in 2 sessions. Movies were collected on an FEI Titan Krios at spot size 5 with a Gatan K3 Camera equipped with an energy filter set to 20 eV. Collection was performed using Leginon software^34^. In both sessions, movies were collected in super-resolution mode at 0.4125 Å/pix at a defocus range of 0.9-1.9 μm, and fractionated into 50 frames of 40 ms each. In the first session, 12,250 micrographs were collected at 30.15 e-/Å^2^/s with a total exposure of 2 seconds for a total accumulated dose of 60.29 e-/Å^2^. In the second session, 23,459 micrographs were collected at 28.97 e-/Å^2^/s with a total exposure of 2 seconds for a total accumulated dose of 57.95 e-/Å^2^ (For full dataset details see Table 1).

#### Preparation of γ-TuRC-capped MT cryo-EM grids

Sample was prepared as described in the section “Solution Pelleting Reaction”. Quantifoil R1.2/1.3 400-square mesh copper grids with a 2 nm continuous carbon layer (Quantifoil) were plasma-treated in a Pelco easiGlow glow discharging system (Ted Pella) for 8 s at 10 mA. Grids were loaded on a Leica EM GP2 plunge freezer (Leica Microsystems) set up with a chamber set to 26°C and 95% humidity. 3 μL of γ-TuRC capped MT sample was applied to the grids with a 200 μL pipette tip with the end cut off (to create a larger diameter to avoid shearing the MTs). The sample incubated in the chamber for 30 seconds before manually blotting. This step was repeated 1 time with a 45 second incubation to apply additional sample. Finally, the grid was washed with 3 μL of BRB80 + 0.025% (v/v) Tween 20 and blotted for 8 seconds by the Leica EM GP2. The blot paper was set up to contact ∼1/2 of the grid to create a gradient of ice. The grid was then immediately plunged into liquid ethane and transferred to liquid nitrogen for storage.

Data was collected at the BioKEM EM Facility at University of Colorado Boulder. Movies were collected on a Titan Krios G3i at spot size 7 with a Falcon 4 Direct Detector (Thermo Fisher Scientific) equipped with an energy filter set to 20 eV. Collection was performed using SerialEM software^35^. Movies were collected in counting mode at 1.663 Å/pix and fractionated into 10 frames per tilt angle of 185 ms each. 41 tilt angles were collected using a continuous strategy from −60° to 60° with a 3° step size. 105 total tilt series were collected with a total dose of 110 e-Å^2^.

### EM data processing

#### Data processing for single particle γ-TuRC with γ-TuNA data

The γ-TuRC single particle data was processed using cryoSPARC v3.0 to v4.2.1^17^. Datasets were imported into cryoSPARC and processed separately up until 3D homogenous refinement (Supplementary Figure 3). Movies were motion corrected using the patch motion correction algorithm, binned by 2 for a working pixel size of 0.825 Å/pix, and the CTFs of the resulting micrographs were estimated using Patch CTF estimation. 3,557 particles were manually picked from 300 micrographs and 878 good particles were selected after 2D classification. These particles and micrographs were then used to train *topaz* pickers^36^, and were then applied to dataset 1 and dataset 2 to pick ∼1.5 million and ∼2.9 million particles respectively. After removing duplicate particles, particles from each dataset were extracted with a box size of 545 Å. 3-4 rounds of 2D classification were performed to select particles and exclude junk. Still keeping the two datasets separate, selected particles from each dataset were used to create 3 ab initio maps of γ-TuRC and particles from rejected 2D classes were used to create 3 “decoy maps” for further classification of particles. We used 2-8 rounds of 3D heterogeneous refinement to sort good particles from bad particles by importing the best ab initio map of γ-TuRC and either two or three “decoy” maps’. Particles sorted into the decoy map were discarded at each iteration until the resolution of the map stopped improving. Dataset 1 contained in 132,189 particles and dataset 2 contained in 162,477. The particles from dataset 2 were refined using an ab initio map generated from these particles as an initial model. This map was then used as a reference for 3D homogenous refinement with all 249,546 particles from both datasets. This resulting map is referred to as the consensus map.

Next, 3D variability analysis^18^ was performed on the consensus map to analyze any large-scale changes in conformation of γ-TuRC. The output was low-pass filtered to 10 Å to observe larger-scale changes, and we set the job up to solve 3 modes. A very loose mask (Supplementary Figure 6, yellow mask) was used to allow for large changes in conformation, as a tight mask may cut off density that varies too much from the consensus structure. The 3D Variability Analysis in simple output mode gave 3 volume series corresponding to 3 components of movement displayed in Supplementary Movies 1-3. In parallel, 3D variability analysis was also run in cluster mode using 4 classes. Two of the four output classes, Class 1 and Class 2, were selected for further processing and refined using a 3D homogenous refinement strategy.

To analyze heterogeneity with cryoDRGN^19^, 294,526 particles and associated poses and CTF were extracted from the consensus map from CryoSPARC. Particles images were downsampled to size 256×256. A cryoDRGN v2.3.0 model was trained for 40 epochs with a latent dimension of 8. All other hyperparameters were left at their default values. The cryoDRGN landscape analysis pipeline was used to generate conformation space plots and volume trajectories. First, the *analyze_landscape* command was used to perform principal component (PC) analysis and hierarchical clustering on 1,000 generated volumes. The mask threshold was set to 0.12 and ward linkage was used as the hierarchical clustering criterion. Then, the *analyze_landscape_full* command was used to embed all particles in the volume PC space. Class labels—either Class 1, Class 2, or neither class—for each particle were extracted from the CryoSPARC 3D variability analysis, and the resulting labels were used to generate contours on the cryoDRGN volume PC plot. The conformation with disordered spoke 11-14 density was validated through a homogenous reconstruction in cryoDRGN.

To improve the local resolution of spoke pairs, masks were created surrounding each spoke pai in Chimera^37^, by erasing the surrounding density, followed by low-pass filtering with a 3 voxel Gaussian filter, and finally thresholding at an appropriate signal-to-noise ratio. Omit masks, for particle subtraction, were also generated by subtracting the spoke pair mask from the unmasked density. These chimera-prepared masks were then imported into cryoSPARC^17^, where a 3 voxel soft edge was applied. Following particle subtraction, local refinement with non-uniform regularization was performed, resulting in locally-refined maps of Class 1 and Class 2. These maps being of insufficient resolution for spokes 11-14, locally-refined maps were created from the full set of particles derived from the consensus map. Next, 3D classification was performed using the following approach: (1) A cryoSPARC 3D-variability job^18^ was run on the locally-refined consensus map; (2) Six volumes were created across the 3 principal components via a 3D variability display job; (3) The two extremes of the volume series along the first principal component (element 1 and element 6) were re-imported into cryoSPARC and used as inputs to a cryoSPARC 3D classification job; (4) The two output classes were separately refined using local refinement with non-uniform regularization as above. For spokes 11-12, one of these two classes resulted in a sub-4 Å resolution map (Table 1), whereas the other consisted of poorly-aligned particles at a resolution of 4.4 Å. However, for spokes 13-14, resolutions remained between 5 and 6 Å.

To further improve the resolution of spokes 13-14, additional rounds of 3D classification were added to the local refinement strategy. Because prior work had suggested spokes 13-14 to be labile^14^, an initial local refinement was performed on spokes 11-14, followed by a single round of 3D classification using as input both the locally-refined spoke 11-14 map, as well as a map where spokes 13-14 were manually erased in Chimera. Based on this classification, 10% of the γ-TuRC particles were most consistent with dissociation of spokes 13-14. The remaining 90% of intact particles were subjected to a second round of 3D variability-dependent 3D classification was performed, in the same manner as for spokes 11-12 above, resulting in a sub-4 Å resolution map (Table 1).

For the purposes of model building and refinement, composite maps for Class 1 and Class 2 were created through superposition of locally-refined maps. The high resolution Class 1 or Class 2 locally-refined maps were confirmed to be in the same position as the overall Class 1 or Class 2 map. The high-resolution maps of the consensus structures of spokes 11-12 and spokes 13-14 were superimposed using *fit in map* upon the respective low-resolution section of the overall Class 1 and 2 maps. Finally, the seven spoke-pair maps were combined using *vop maximum*.

#### Data processing for γ-TuRC capped MT cryo-ET data

Tilt series collected on the Krios were motion corrected, aligned and reconstructed into tomograms as detailed below using IMOD v.4.12.3^38, 39^. IMOD *alignframes* was used to motion correct each of the 41 movies in each tilt series and create an ordered (−60° to +60°) tilt series of the resulting micrographs. Each tilt series was used as input into the IMOD *etomo* pipeline to create tomograms using the batch processing method. Patch tracking was used to align stacks, as no fiducials were added to the sample. Each tomogram was individually checked to ensure quality of processing. Tomograms were binned by 4 in IMOD to increase contrast before particle averaging.

Next, individual γ-TuRC capped ends were picked from each tomogram and averaged using PEET v.1.15.0^40, 41^, as detailed below. 2-point models were created using *3dmod* with the head point being placed at the base of γ-TuRC and the tail point being placed down the center of the MT axis. IMOD *stalkInit* was then used to create a model centered on γ-TuRC as well as an initial particle alignment relative to the MT axis. The model and alignment files were then input to PEET. 94 total sub-volumes were iteratively refined and averaged to create the final sub-tomogram average. A mask covering the MT capped end was created in USCF Chimera and binarized, padded, and given a soft edge in Relion 3.1.0. Tomograms were then split into even and odd sets, and half-maps were averaged and calculated separately. These half-maps were input into the cryoSPARC *FSC* utility and plotted. Because the estimated 34 Å resolution is limited by the low number of sub-tomograms and preferred orientations within the tomograms, the results were appraised solely at a domain level. Maps were low-pass filtered to an appropriate resolution for analysis, typically 34 Å using cryoSPARC. Class 1 and Class 2 structures were manually placed into the density, then rigid-body fit into the reconstruction using *fit in map* in USCF Chimera.

### Model building and analysis

#### Model building for γ-TuRC with γ-TuNA

Molecular models of *Xenopus laevis* γ-TuRC was determined by first placing the previously-reported *X. laevis* γ-TuRC structure, determined at 4.8 Å resolution (PDB ID 6TF9^11^), into the homogeneously-refined density maps for both Class 1 and Class 2 and the overall fit adjusted by rigid-body refinement using the real_space_refine program of the Phenix suite^42^. An improved fit was obtained by rigid-body refinement treating individual domains within each GCP and γ-tubulin separately. Inspection of the fit to the density maps revealed a number of interpretation discrepancies within each GCP for this starting model. Models for γ-TuRC were extensively re-fit into the composite Class 1 and Class 2 maps, with manual rebuilding and local real-space refinement using the program COOT^43^. The model for the lumenal belt was taken from the human γ-TuRC structure (PDB ID 6V6S, resolution 4.3 Å^10^), mutated to match the *X. laevis* sequence, and extended with additional polypeptide chain for GCP6 and GCP3. The staple peptide at the N-terminus of GCP2 was taken from the initial human backbone trace which was updated and then interpreted in terms of the *X. laevis* sequence. Additional extended polypeptide chain identified as belonging to GCP6 was found on the inside of the structure extending from the GCP2 in the first spoke through the lumenal belt and GCP6 in spoke 12. Four short α-helices that are otherwise unassigned to a specific protein span the inside surface of the GCPs from GCP6 in spoke 12 through GCP3 in spoke 14. After rebuilding and local real space refinement, the Class 1 and Class 2 models were subject to real-space refinement of the entire structure using *real_space_refine* at a nominal resolution of 3.0 Å. Refinement statistics can be found in Table 1. In accordance with prior convention^11^, the sequences of L-chromosome homeologs were used for building when two homeologs were present,.

#### Analysis of γ-TuRC Structure

To calculate the motion of γ-tubulin molecules between Class 1 and Class 2, the position of each γ-tubulin was recorded for the two conformers, which are already globally aligned to one another and the consensus map. Next, each γ-tubulin for Class 1 was superimposed onto the analogous γ-tubulin in Class 2 using Cα superposition in PyMOL^44^. Superposition vectors were determined by measuring the displacement of the Cα of Tyr-169, which marks the approximate centroid of the molecule, between the global superposition and the local superposition. The components of the displacement vector were then extracted in a reference frame composed of three vectors for the Class 1 γ-tubulin: Ile-318 Cα to Arg-124 Cα (left-right), Arg-244 Cα to Asn-102 Cα (down-up), Ser-226 Cα to Asp-159 Cα (in-out). Displacement vectors were visualized using the PyMOL add-on *modevectors* authored by Sean M. Law.

Helical parameters for the Tyr-169 Cα centroids of the γ-tubulin helix were determined using the *Define Axes* program in UCSF Chimera with helical correction, and then distances were computed between the Cα of each Tyr-169 and the helical axis.

Buried surface area was calculated using the PISA server^45^ with the following domain assignments: for GCP2, GRIP1 comprises residues Glu-213 through Glu-504 and GRIP2 Leu-507 through Glu-869; for GCP3, GRIP1 comprises residues Glu-244 through Leu-546 and GRIP2 Leu-552 through Val-892; for GCP4, GRIP1 comprises residues Ile-2 through Glu-347 and GRIP2 Leu-350 through Tyr-654; for GCP5, GRIP1 comprises residues Glu-263 through Asp-708 and GRIP2 Leu-711 through Leu-1012; for GCP6, GRIP1 comprises Glu-349 through Val-1387 and GRIP2 Met-1391 through Tyr-1594. Domain assignments were computed by a combination of sequence alignment and secondary structure assignment. Surface areas were computed based on the domain assignments for the left-most spoke, such that contacts between the γ-tubulin of spoke *n* on the GRIP2 domain of spoke *n+1* were accounted as γ-tubulin contacts.

The distance between adjacent γ-tubulins was calculated from the distance between adjacent γ-tubulin residue Tyr-169 Cαs, for yeast γ-tubulin, the homologous residue Tyr-170 Cα, and for the MT, the β-tubulin homologous residue Phe-167 Cα. The angle adopted by GCP spokes was calculated as described in^11^.

## ACKNOWLEDGEMENTS

Screening was carried out at the Princeton IAC and single-particle cryo-EM data collection was carried out at the NYU cryo-EM core facility. Cryo-electron tomography data was collected at the CCET through the NIH Tomography Network. We thank Paul Shao and John Schreiber for support in the Princeton IAC, Bill Rice and Bing Wang for support in the NYU cryo-EM facility, Courtney Ozzello, Garry Morgan, John Heumann, and Andreas Hoenger from CCET, and Charles Moe from the BioKEM Facility at UC Boulder. We also thank Venecia Valdez for preparing reagents used in this study, Matthew Cahn for computational support, Wei Dai and Jason Kaelber for their support and guidance on the tomography project, Jonathan Bouvette for technical advice, Frederick Hughson for scientific discussions and critical reading of the manuscript, and Damian Ekiert for guidance on model building and validation.

This work was supported by National Institutes of Health grants F32GM142149-01A1 (SMT), NIGMS grant R01 1R01GM141100-01A1 (SP), NIGMS grant R00GM112982 (GB), Damon Runyon Cancer Research Foundation DFS-20-16 (G.B.). The authors acknowledge the use of Princeton’s Imaging and Analysis Center (IAC), which is partially supported by the Princeton Center for Complex Materials (PCCM), a National Science Foundation (NSF) Materials Research Science and Engineering Center (MRSEC; DMR-2011750). Some of the work was performed at NYU Langone Health’s Cryo-Electron Microscopy Laboratory (RRID: SCR_019202), which is partially supported by the Laura and Isaac Perlmutter Cancer Center Support Grant NIH/NCI P30CA016087. Molecular graphics and analysis was performed with UCSF Chimera, developed by the Resource for Biocomputing, Visualization, and Informatics at the University of California, San Francisco, with support from NIH P41-GM103311.

## DATA AVAILABILITY

The reconstructed density maps will be submitted to and accessible from the EMDB. The structural models of Class 1 and Class 2 will be submitted to and accessible from the PDB.

## AUTHOR CONTRIBUTIONS

Brianna Romer, Brian P. Mahon, and Michael J. Rale conceived of the project. Sabine Petry and Gira Bhabha supervised the project and acquired funding. Brianna Romer, Brian P. Mahon, and Michael J. Rale generated samples. Brianna Romer and Brian P. Mahon made cryo-EM grids and collected cryo-EM data. Brianna Romer and Sophie M. Travis processed cryo-EM data with guidance from Nicolas Coudray. Philip D. Jeffrey built, refined, and validated the structural models with assistance from Sophie M. Travis and Collin T. McManus. Rishwanth Raghu and Ellen D. Zhong performed and interpreted the cryoDRGN analysis. Sophie M. Travis performed model analysis. Collin T. McManus and Brianna Romer collected and analyzed TIRF data. Sophie M. Travis and Brianna Romer wrote the manuscript. All authors edited the manuscript.

