## Supplementary Information for "Conformational states of the microtubule nucleator, the γ-tubulin ring complex"

for

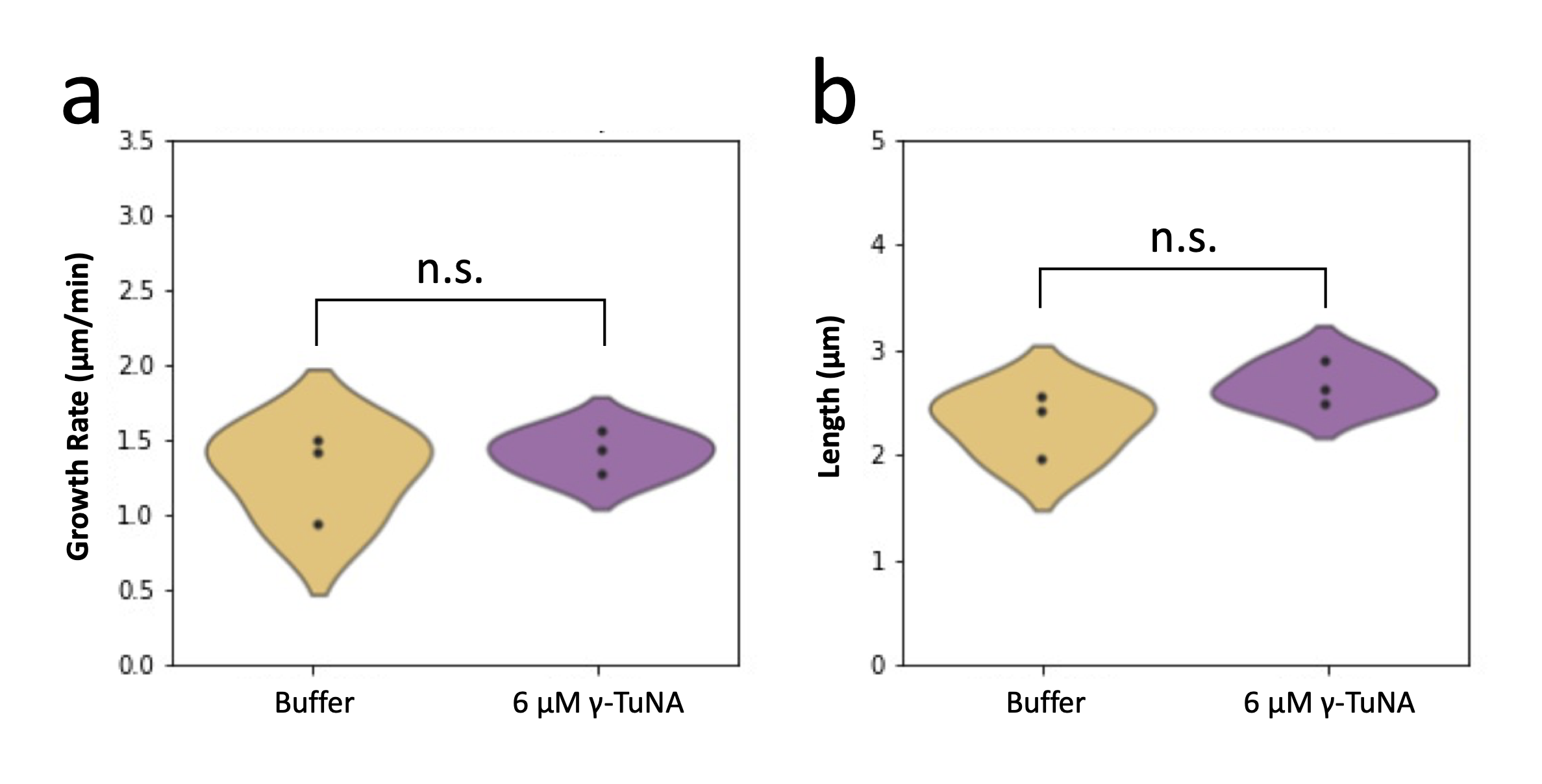

**Supplementary Figure 1: Other reaction properties of MT nucleation by endogenous purified *Xenopus laevis* γ-TuRC**

a. Quantification of the mean growth rate of γ-TuRC nucleated MTs. Unactivated γ-TuRC nucleated MTs (yellow) grew at an mean rate of 1.42 μm/s, and activated γ-TuRC MTs (purple) grew at an average rate of 1.29 μm/s. Rates are calculated for the 3 independent reactions also shown in Figure 1b.

b. Quantification of the maximum length achieved for each γ-TuRC nucleated MT. Unactivated γ-TuRC nucleated MTs (yellow) had an average length of 2.31 μm, and activated γ-TuRC nucleated MTs (purple) had an average length of 2.67 μm. Maximum lengths are calculated for the 3 independent reactions also shown in Figure 1b.

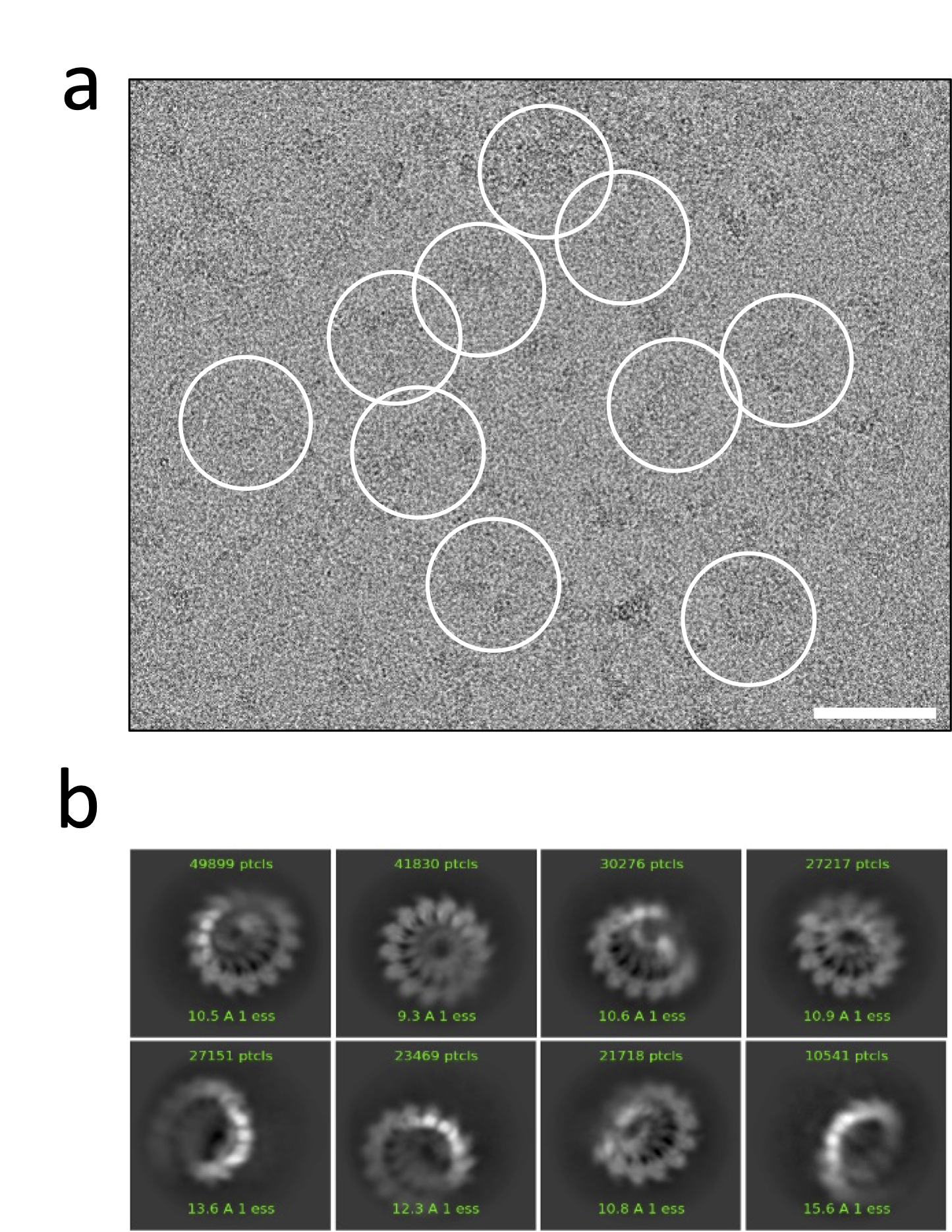

**Supplementary Figure 2: *Xenopus laevis* γ-TuRC cryo-EM raw data**

a. A representative micrograph showing picked γ-TuRC particles within white circles. Scale bar is 50 nm and circle diameter is 660 pixels (545 A).

b. Representative 2D class averages of γ-TuRC particles. Each box side is 545 Å.

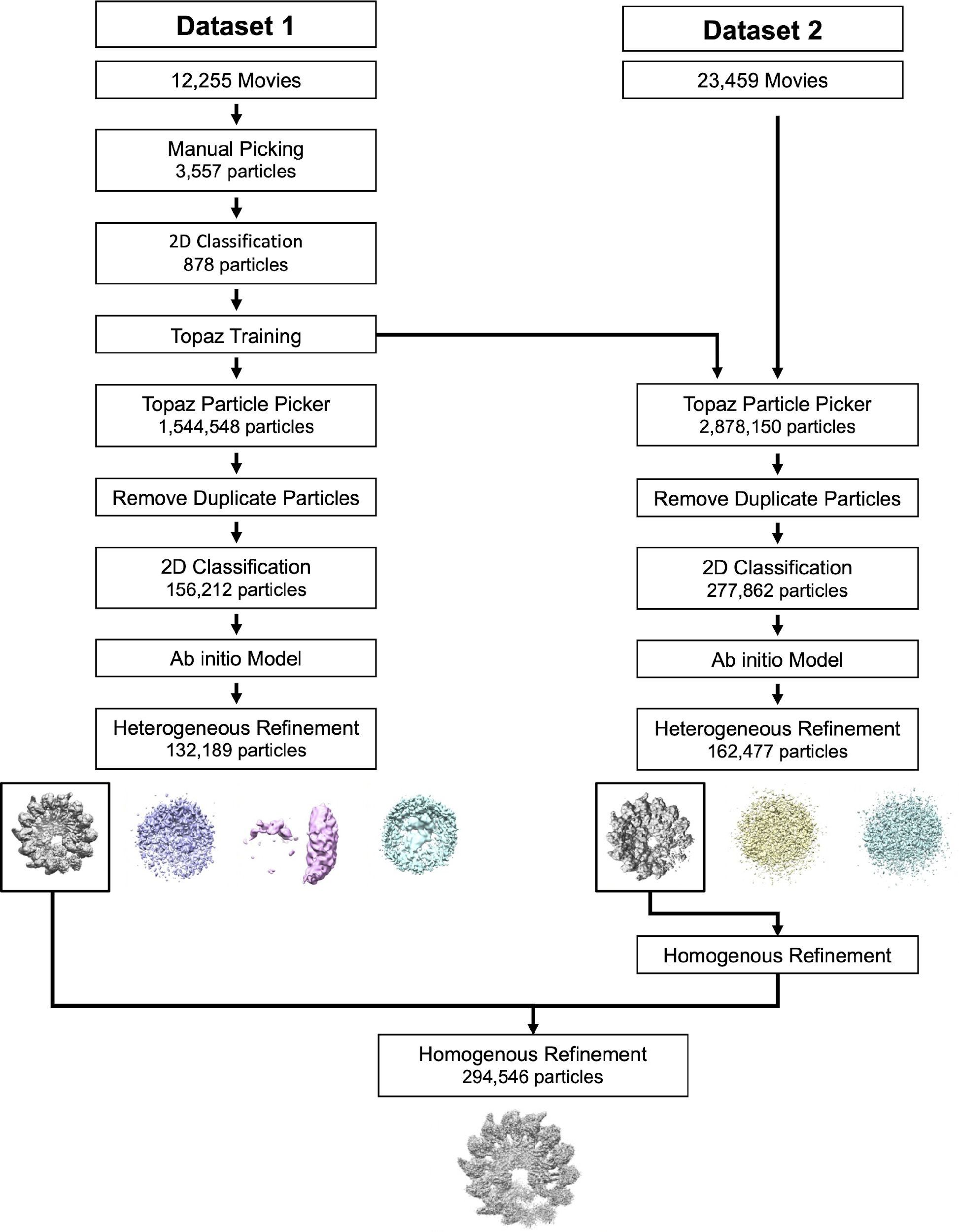

**Supplementary Figure 3: Cryo-EM data processing workflow for consensus map**

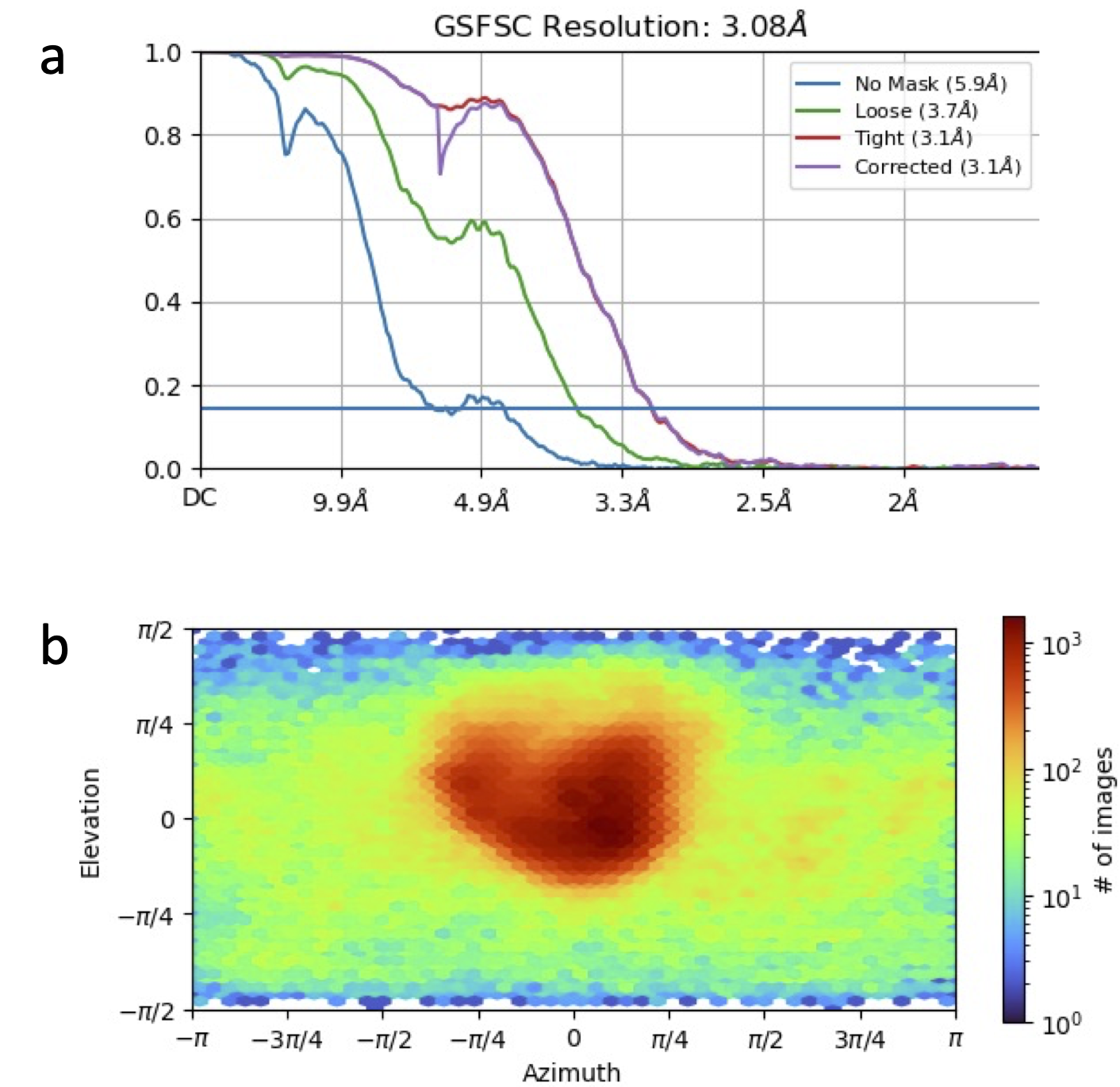

**Supplementary Figure 4: Consensus map properties**

a. Resolution estimate via FCS curve

b. Distribution of particles by viewing direction

**Supplementary Figure 5: Properties of the Class 1 and Class 2 maps**
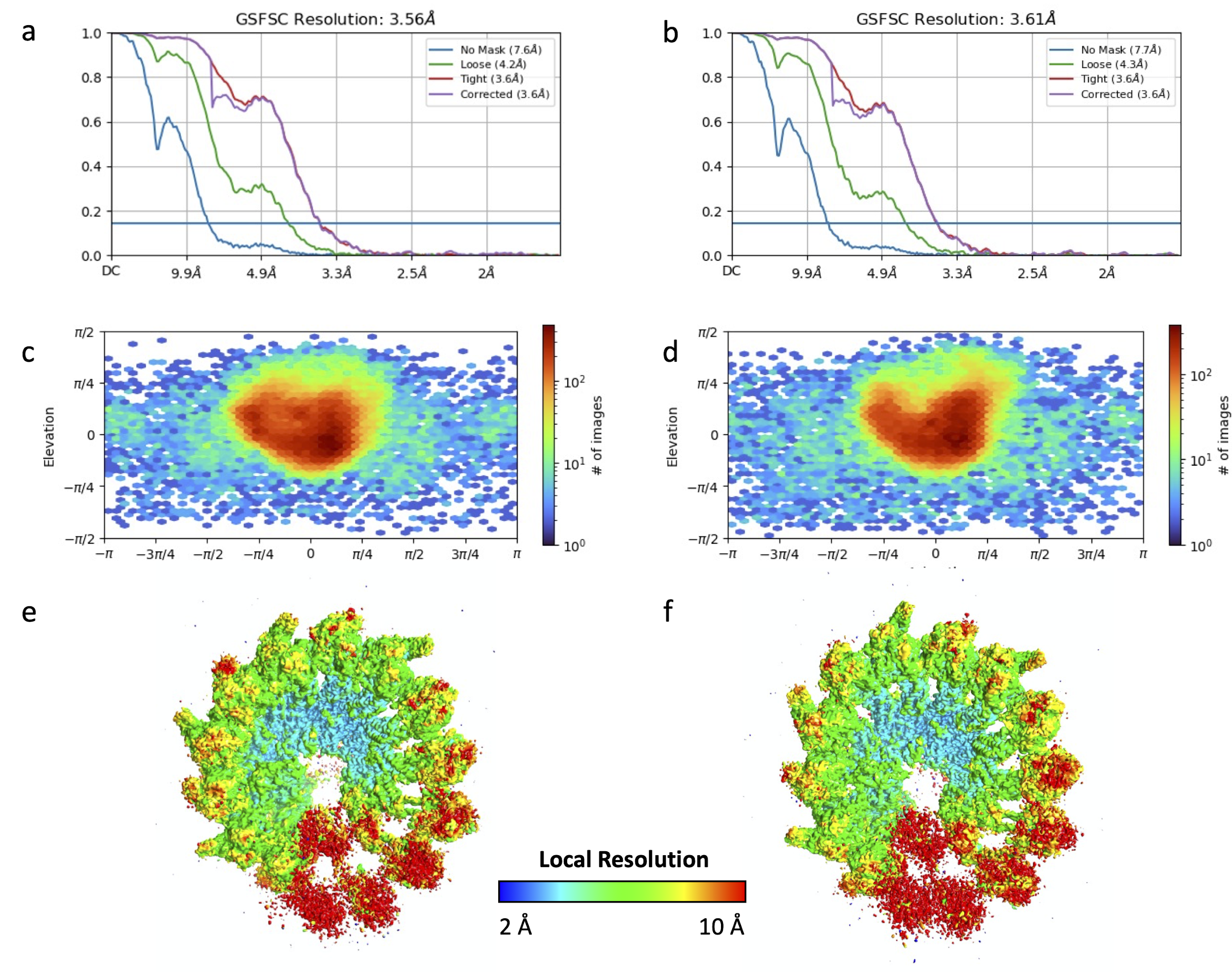

a-b. Resolution estimate via FCS curve for Class 1 (a) and Class 2 (b)

e-d. Distribution of particles by viewing direction for Class 1 (c) and Class 2 (d)

e-f. Cryo-EM density map colored by local resolution for Class 1 (e) and Class 2 (f), where resolution ranges from 2 Å (dark blue) to 10 Å (red).

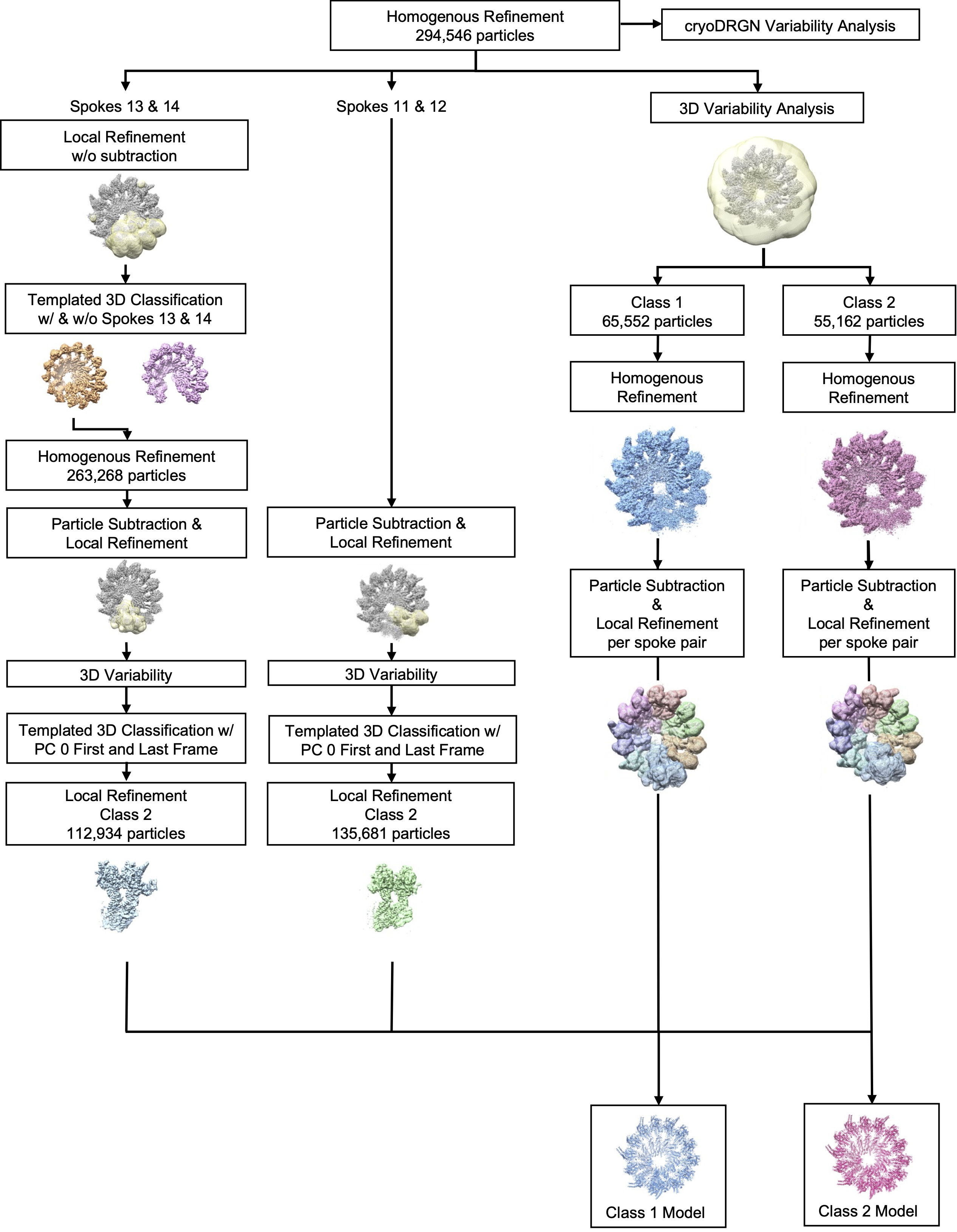

**Supplementary Figure 6: Cryo-EM data processing workflow for local refinement of γ-TuRC spoke pairs**

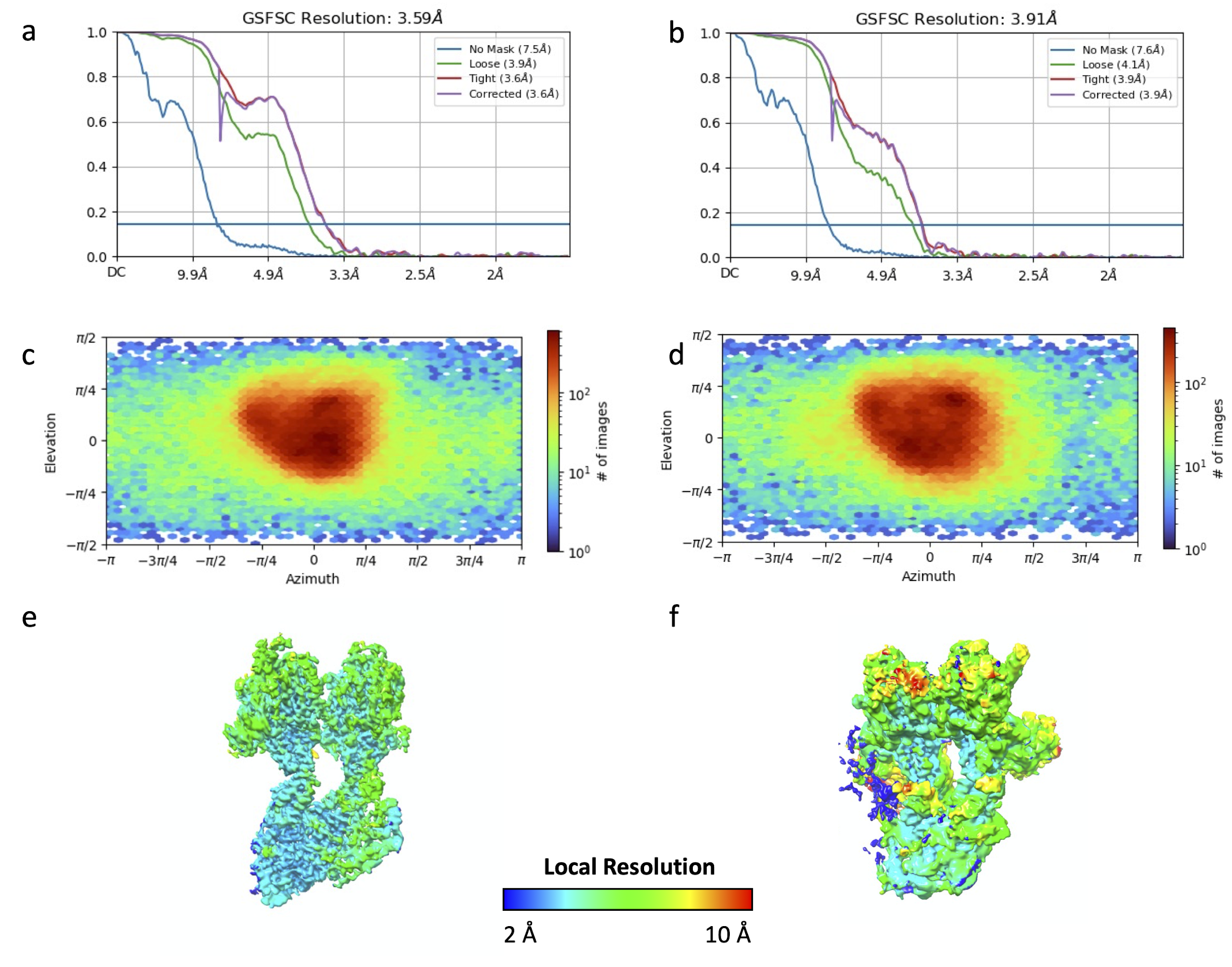

**Supplementary Figure 7: Properties of the spoke 11-12 and spoke 13-14 maps**

a-b. Resolution estimate via FCS curve for spoke 11-12 (a) and spoke 13-14 (b)

c-d. Distribution of particles by viewing direction for spoke 11-12 (c) and spoke 13-14 (d)

e-f. Cryo-EM density map colored by local resolution for spoke 11-12 (e) and spoke 13-14 (f), where resolution ranges from 2 Å (dark blue) to 10 Å (red).

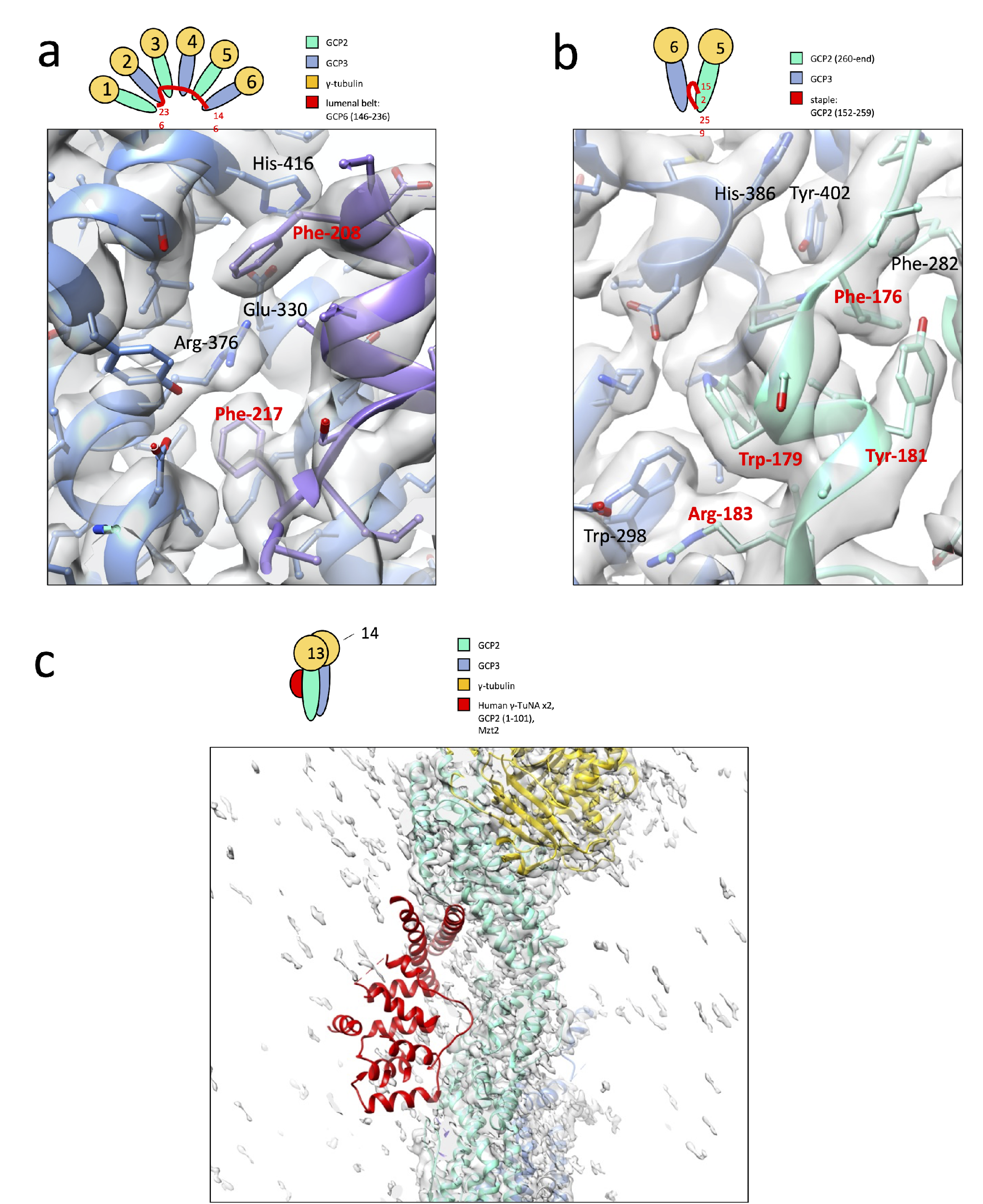

**Supplementary Figure 8: Model fit into cryo-EM density map of selected γ-TuRC regions**

a. Above, cartoon showing the location of the lumenal belt (GCP6 146-236, red) bound to the lumenal portions of spokes 2-6. Below, model of a portion of the lumenal belt (purple model, red residue labels) at spoke 2 showing the docking of bulky aromatic residues into the cryo-EM density map (gray surface). Nearby and contacting residues of the GRIP1 domain of spoke 3 GCP3 are modeled in blue. Models and maps shown are for Class 1 here and in the remainder of the figure.

b. Above, cartoon showing the location of the staple peptide (GCP2 152-259, red) on the outside of γ-TuRC between the GCP2 and GCP3 subunits of a γ-TuSC. Below, model of a portion of the staple peptide (cyan model, red residue labels) between spokes 5 and 6 showing the docking of bulky aromatic residues into the cryo-EM density map (gray surface). The staple peptide makes contact with aromatic residues of the GRIP1 domains of both GCP2 (cyan model, black labels) and GCP3 (blue model).

c. Above, cartoon showing the location of the γ-TuNA dimer bound to the N-terminus of GCP2 and Mzt2 (red) on the outside of spoke 13 as in the human γ-TuRC structure (PDB accession 6V6S, remodeled as 6X0Z). Below, alignment of human γ-TuNA dimer, N-terminus of GCP2, and Mzt2 (red, PDB accession 6X0Z) on the outside of the GCP2 of spoke 13 (cyan model), showing that, in our cryo-EM density map (gray surface), no density is present for this feature.

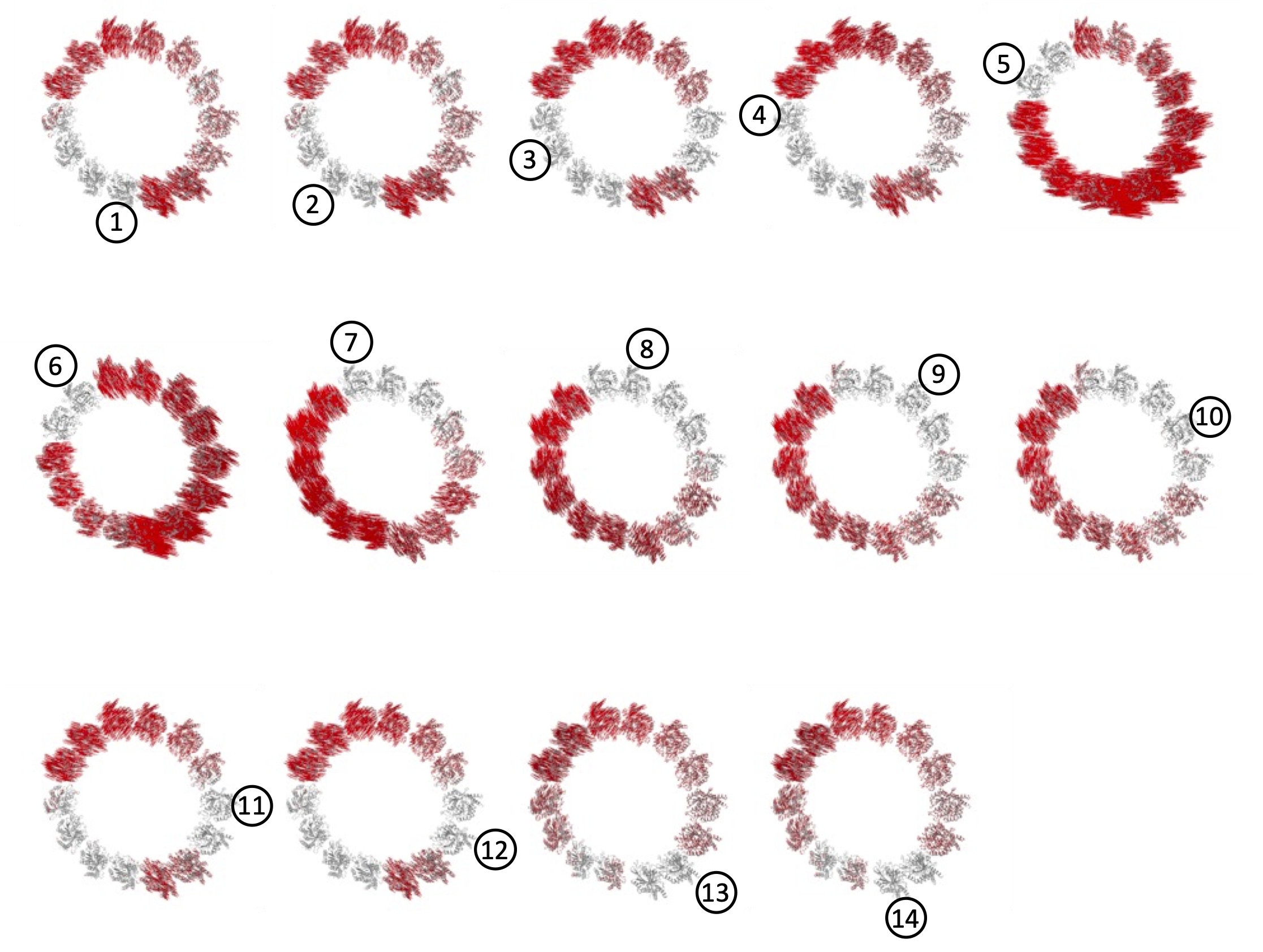

**Supplementary Figure 9: Spoke-centric displacement diagrams between Class 1 and Class 2**

γ-tubulins of individual spokes (indicated spoke shown as a circled number) were superimposed from Class 2 onto Class 1. Then, displacement vectors were calculated and displayed for all γ-tubulin Cαs from Class 1 to Class 2. Displacement vectors are displayed as red lines.

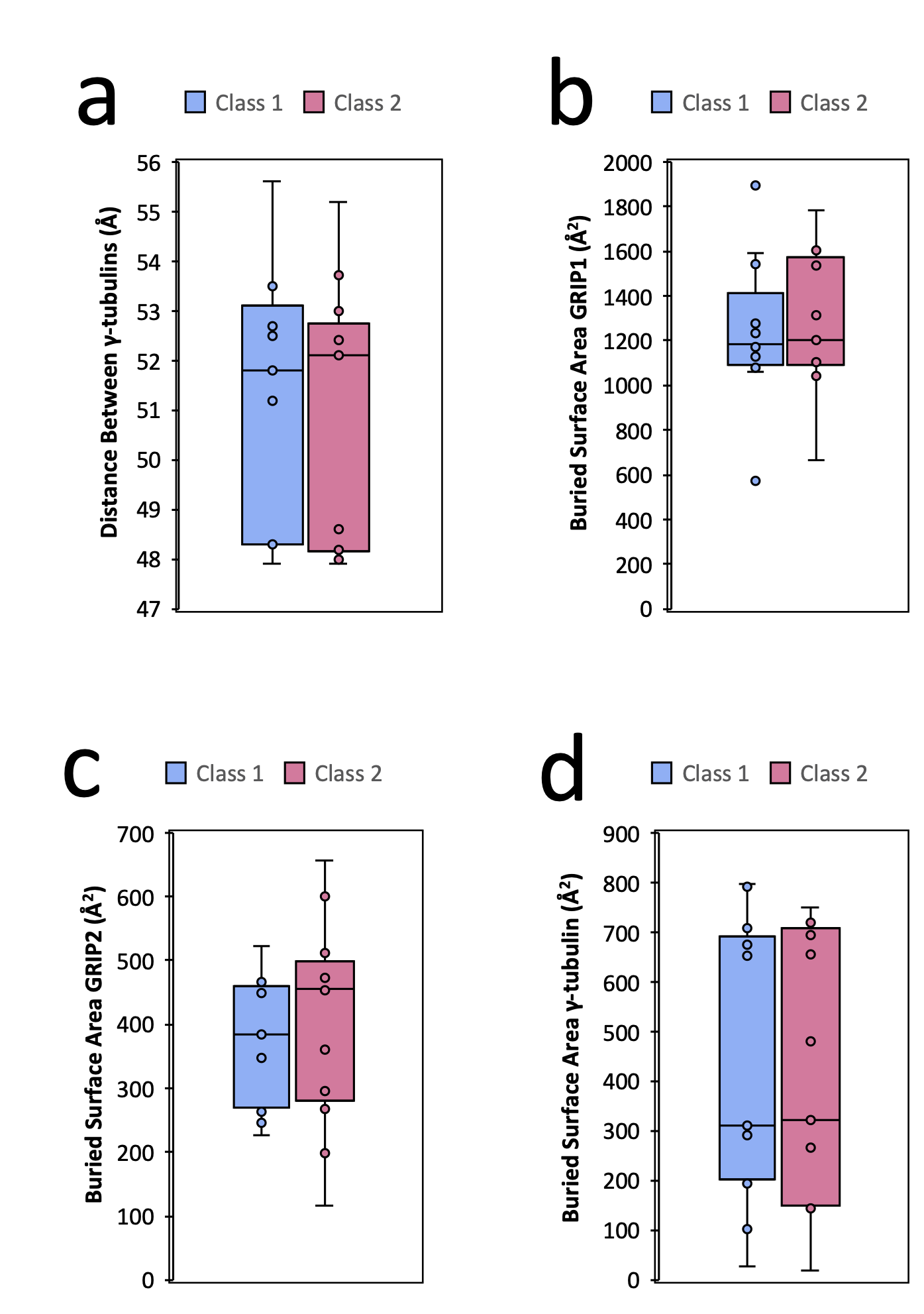

**Supplementary Figure 10: Statistical analysis of structural properties comparing Class 1 vs. Class 2**

a. Further analysis of γ-tubulin distance data presented in Figure 3b. Median shown as central line, and middle two quartiles shown in colored box.

b. Further analysis of GRIP1 buried surface area data presented in Figure 3d

c. Further analysis of GRIP2 buried surface area data presented in Figure 3d

d. Further analysis of γ-tubulin buried surface area data presented in Figure 3d

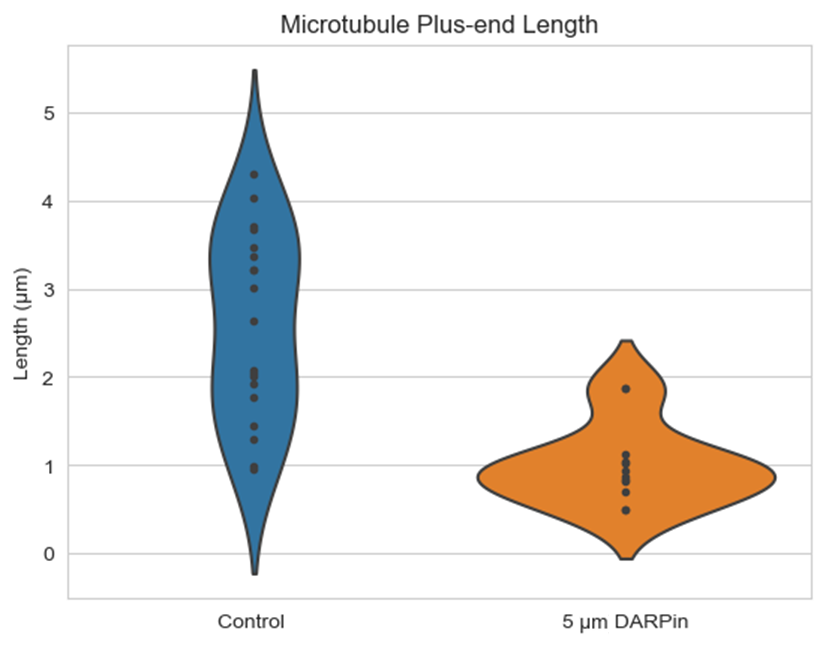

**Supplementary Figure 11: Quantification of length of plus-ends of polymerized MTs**

Polymerization of MT lattice from stabilized MT seeds was quantified in the presence or absence of 5 μM DARPin TM-3 after 5 minutes. N=19 for control, N=12 for 5 μM DARPin TM-3.

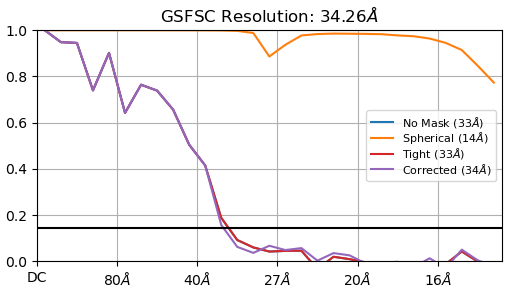

**Supplementary Figure 12: FSC curve for sub-tomogram average of capped MT end**

After randomly splitting the subtomogram volumes into two half-data sets, volumes were re-aligned in PEET and FSC was calculated using cryoSPARC.

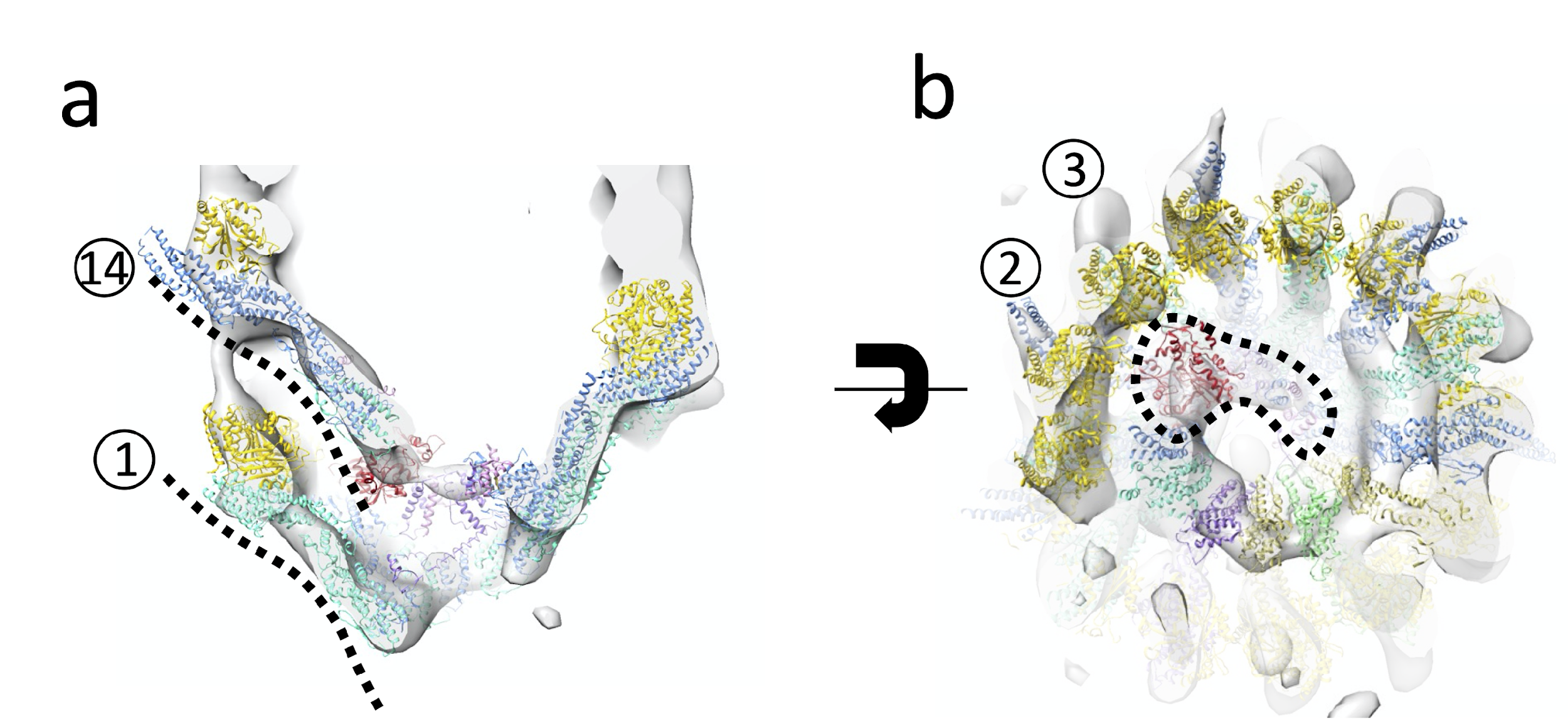

**Supplementary Figure 13: Asymmetric features of the capped MT end supporting placement of γ-TuRC**

a. Overlap of spokes 1 and 14. The contour of spokes 1 and 14 are outlined with a dashed black line.

b. Black dashed outline highlighting density for luminal bridge, most prominently the actin monomer found next to the base of spokes 2 and 3.

**Supplementary Figure 14: Local ordering of spokes 13 and 14 between apo and post-nucleation γ-TuRC**
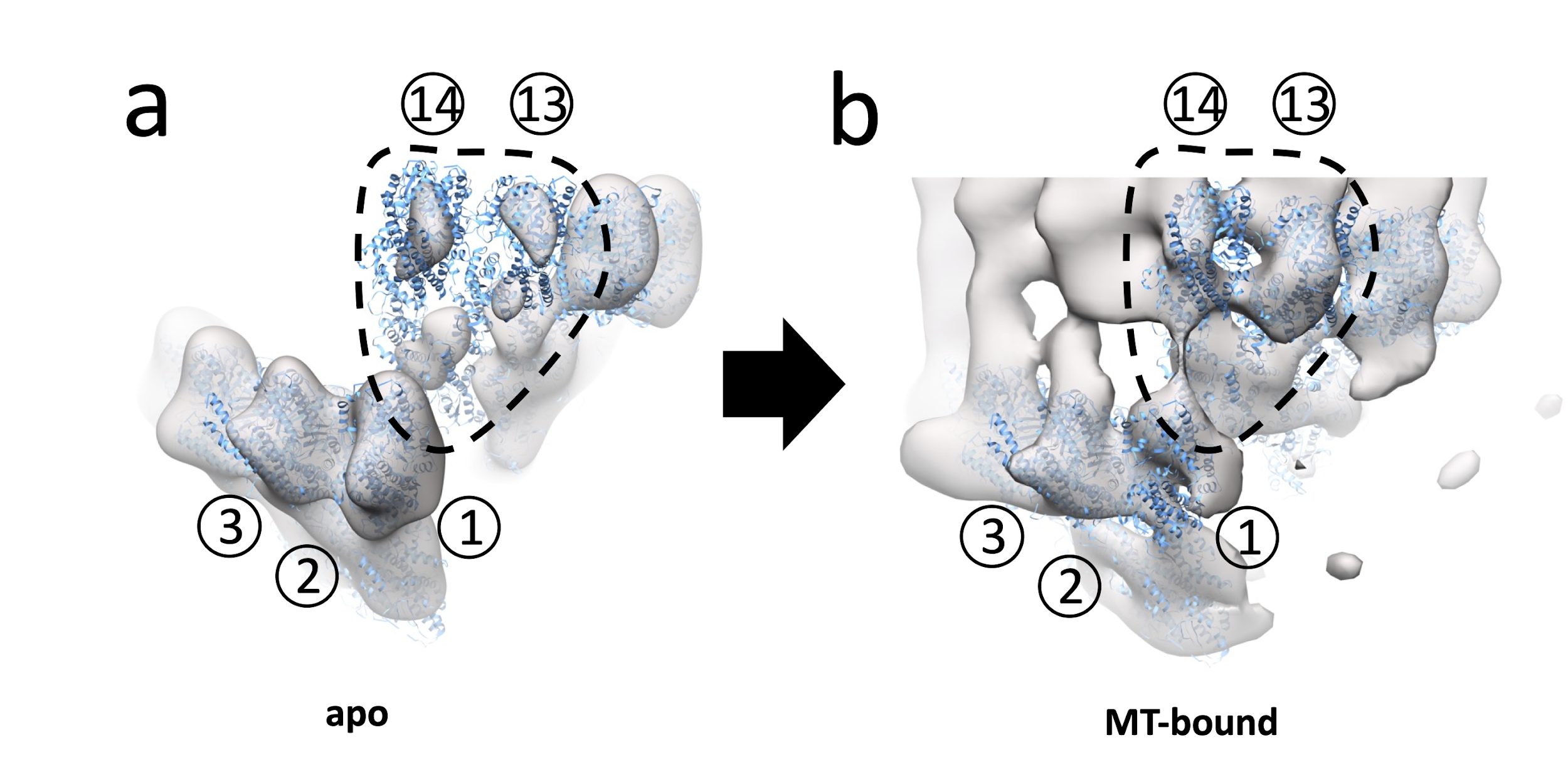

a. Map of Class 1 γ-TuRC, low-pass filtered at 34 Å to match the cryo-ET reconstruction, shows a substantial lack of density for spokes 13 and 14 when fit to the Class 1 model.

b. Cryo-ET reconstruction of the post-nucleation state of γ-TuRC, shown in the same orientation, shows approximately the same amount of density for spokes 13-14 as for spokes 1-2. In both panels, maps are shown as translucent gray surfaces and the structure of Class 1 γ-TuRC as a blue cartoon. Thresholds were chosen such as the volume of spoke 1 is visually roughly similar in both maps. Selected spokes are numbered and spokes 13-14 are circled with a dashed black line.

**Table 1: Cryo-EM data collection, refinement, and validation statistics**

|  | Class 1 overall  (EMDB-TBD) | Class 2 overall  (EMDB-TBD) | Spoke 11-12 (EMDB-TBD) | Spoke 13-14 (EMDB-TBD) | Class 1 composite  (EMDB-TBD)  (PDB-TBD) | Class 2 composite  (EMDB-TBD)  (PDB-TBD) | Capped MT (EMDB-TBD) |
| --- | --- | --- | --- | --- | --- | --- | --- |
| **Data collection** |  | | | |  |  |  |
| ***Titan Krios settings*** |  | | | |  |  |  |
| Voltage (kV) | 300 | | | |  |  | 300 |
| Defocus range (μm) | -0.9 to -1.9 | | | |  |  | -2 to -4 |
| Energy filter width (μm) | 20 | | | |  |  | 20 |
| ***Detector settings*** |  | | | |  |  |  |
| Detector type | Gatan K3 | | | |  |  | Falcon 4 |
| Exposure time (ms) | 2000 | | | |  |  | 699 |
| Magnification | 105 kx | | | |  |  | 13 kx |
| Acquisition mode | Super-resolution | | | |  |  | Counting |
| Pixel size (Å) | 0.4125 | | | |  |  | 1.663 |
|  | **Dataset 1** | | **Dataset 2** | |  |  |  |
| Dose rate (e^-^/Å^2^/s) | 30.15 | | 28.97 | |  |  | 1.448 |
| Total dose (e^-^/Å^2^) | 60.29 | | 57.95 | |  | Tilt Dose | 2.683 |
| Total frames | 50 | | 50 | |  | Tilt frames | 10 |
| Number of Movies | 12,255 | | 23,459 | |  |  |  |
| ***Tomography settings*** |  |  |  |  |  |  |  |
| Number of tilts |  |  |  |  |  |  | 41 |
| Tilt angle step |  |  |  |  |  |  | 3° |
| Tilt angle range |  |  |  |  |  |  | -60° to 60° |
| Tilt strategy |  |  |  |  |  |  | continuous |
| Total dose (e^-^/Å^2^) |  |  |  |  |  |  | 110 |
| Number of tilt series |  |  |  |  |  |  | 105 |
| **Data processing** |  |  |  |  |  |  |  |
| Symmetry imposed | C1 | C1 | C1 | C1 |  |  | C1 |
| Initial particle images (no.) | 4,422,698 | 4,422,698 | 4,422,698 | 4,422,698 |  |  | 94 |
| Final particle images (no.) | 65,552 | 55,162 | 135,681 | 112,934 |  |  | 94 |
| Map resolution (Å)  FSC threshold | 3.6  0.143 | 3.6  0.143 | 3.6  0.143 | 3.9  0.143 |  |  | 35  0.143 |
| Map resolution range (Å) | 3.6-10 | 3.6-10 | 3.6-10 | 3.9-10 |  |  |  |
| **Refinement** |  |  |  |  |  |  |  |
| Initial model used (PDB code) |  |  |  |  | 6TF9, 6V6S | 6TF9, 6V6S | TBD, 7SJ7 |
| Model resolution (Å)  FSC threshold |  |  |  |  | 3.6  0.143 | 3.6  0.143 |  |
| Model resolution range (Å) |  |  |  |  | 500-3.6 | 500-3.6 |  |
| Map sharpening *B* factor (Å^2^) |  |  |  |  | 0.0 | 0.0 |  |
| Model composition  Non-hydrogen atoms  Protein residues  Ligands |  |  |  |  | 123019  15259  15 | 123098  15272  15 |  |
| *B* factors (Å^2^)  Protein  Ligand |  |  |  |  | 70.9  87.6 | 74.8  86.4 |  |
| R.m.s. deviations  Bond lengths (Å)  R.m.s.z.  Bond angles (°)  R.m.s.z. |  |  |  |  | 0.005  Z=0.205  0.827  Z=0.448 | 0.004  Z=0.169  0.734  Z=0.389 |  |
| **Validation**  MolProbity score  EMRinger score  Clashscore  Poor rotamers (%)  C-beta outliers (%) |  |  |  |  | 2.4  1.26  22.4  0.5  0.0 | 2.2  1.30  18.4  0.7  0.01 |  |
| **Ramachandran plot**  Favored (%)  Allowed (%)  Disallowed (%) |  |  |  |  | 89.99  9.65  0.36 | 92.39  7.36  0.25 |  |
| **RAMA-Z**  Whole  Helix  Sheet  Loop |  |  |  |  | -2.81 (0.07)  -1.18 (0.05)  -2.50 (0.14)  -2.70 (0.08) | -1.50 (0.07)  -0.15 (0.06)  -2.24 (0.14)  -2.05 (0.09) |  |
| **Map correlation coefficients**  CC_mask  CC_volume  CC_peaks  CC_box |  |  |  |  | 0.7198  0.7562  0.5391  0.6293 | 0.7206  0.7285  0.5439  0.6443 |  |
